# Mathematical and ecological traits of above and belowground biomass production of beech (*Fagus silvatica* L.) trees growing in pure even-aged stands in north-east France

**DOI:** 10.1101/300210

**Authors:** Noël Le Goff, Jean-Marc Ottorini

## Abstract

Tree biomass and biomass increment equations were specially developed in 1996–1997 to study the ecophysiological functioning of an experimental European beech stand, aged about 30 year-old, in the Hesse forest (NE France). In order to extend such a study to beech stands of different age classes, it was necessary to build biomass and biomass increment equations that could be used for any age; we call them generalized biomass equations.

To build such generalized equations, trees were sampled in different forest plots covering the whole age range. Moreover, in each plot, several trees were chosen to represent the different crown classes (from dominant to suppressed). Sampled trees were felled down and the root system excavated for a sub-sample of trees, for biomass analysis by separating the main compartments of the above and belowground tree parts. When it was not possible to measure the total biomass of a given tree compartment (large trees), wood samples were taken in the concerned compartment. Moreover, equations were built to estimate the biomass of the missing parts of the root system and branch compartments that were likely to have suffered losses during root excavation and tree felling, respectively. Multivariate linear and non-linear models including possible random effects were tested to represent the biomass and biomass increment variations of each tree compartment and of their aggregation in the above and belowground parts of the tree.

Compatible biomass and biomass increment equations for the different tree compartments and their combination in above and belowground tree parts were developed and fitted, allowing the analysis of the variations of the biomass distribution and allocation with tree age. Stem growth efficiency was also calculated and appeared dependent on tree age and tree social status.

The biomass and biomass increment equations established for beech in this study allow the estimation of the biomass and carbon stocks and fluxes (NPP) for the even-aged beech stands of the Hesse forest, whatever the age of the stand; they could also be used to analyze the effects of different silvicultural treatments on the biomass and carbon stocks and fluxes of beech stands, using the available stand growth and yield models developed for beech in France.

## 1. Introduction

Tree samples were analyzed in Hesse forest (NE France) for biomass and biomass increment in years 1996 and 1997, and then 2001 and 2003, in order to estimate the stand biomass and biomass increment of two beech plots installed to study the functioning of the beech ecosystem (Le Goff & Ottorini, 2001; Le Goff & Ottorini, 1998). In these plots, the yearly carbon fluxes were more particularly evaluated since 1996, using the eddy-covariance method (Granier & al, 2000). The biomass and biomass increment equations developed at tree level allowed the comparison of the net primary productivity (NPP) estimated from the yearly CO_2_ fluxes measured with the current stand biomass increment estimated for the appropriate years, using stand inventories: the good agreement observed between the two estimations of the stand NPP made confident with the CO_2_ fluxes measured (Granier & al, 2000).

Specific biomass equations linking the biomass and the biomass increment of above and belowground tree compartments to tree diameter were developed for each of the two experimental plots in Hesse forest, with mean-range ages of 20 and 30 years, respectively (Le Goff & Ottorini, 1998; Le Goff & Ottorini, 2001; Le Goff, 2001 unpublished data). Considering that the ecophysiological studies, especially the analysis of the links between tree growth and environmental factors, conducted in this forest could be extended to younger or older stands, it would be of interest to develop generalized biomass equations that could apply to all the age range of the classical beech rotation (0–120 years), avoiding the need to build new specific biomass equations. Such models were successfully developed earlier by introducing different tree characteristics in the biomass equations, and in particular tree height (*ht*) often combined with tree diameter^1^ (*d*) in the form of *d*^2^ *ht*, or more generally *d^α^ht^β^* (Genet & al, 2011a, b; Shaiek & al, 2011; McElligott & Bragg, 2013; Sileshi, 2014; Zheng & al, 2015).

The biomass and biomass increment equations previously established in Hesse forest were fitted independently for each tree compartment and for the aboveground and belowground compartments, each one taken as a whole. Thus, the constraint of additivity for the tree compartments composing either the aerial or the belowground tree parts was not considered at that time, although it would have been desirable (Repola, 2008, 2009; Genet & al, 2011a; Parresol, 2011; Zheng & al, 2015).

The main objective of this study was then to develop generalized and compatible biomass and biomass increment equations for beech in the Hesse forest, for the above and belowground tree compartments. Moreover, the study aimed at analyzing the contribution of the different above and belowground tree compartments to tree biomass (biomass distribution) and to tree biomass increment (biomass allocation)^2^.

Prediction models for leaf biomass (and leaf area) will also be considered, and used to analyze the stem growth efficiency (*GE*) of beech trees, *GE* being defined as stem growth per unit of leaf area. This concept of growth efficiency is widely used to identify the silviculturally important patterns of tree and stand productivity (DeRose & Seymour, 2009; Hofmeyer & al, 2010; Konôpka & al, 2010; Maguire & al, 1998; Seymour & Kenefic, 2002), and the variations of *GE* with the growth related tree characteristics (age, leaf area, canopy position) will be examined more particularly in this study for beech.

## 2. Material and Methods

### 2.1 Study site

The study was conducted in the state forest of Hesse, located in the East of France, about 80 km east of Nancy (48°40’ N, 7°05’ E; altitude: between 270 and 330 m), and covering an area of around 450 ha. It is a high forest, naturally regenerated, and composed mainly of oak (*Quercus petraea* and *Q. robur*, 40%) and European beech (*Fagus silvatica* L., 37%), the remaining species being mostly hardwoods (23%). The climate is continental with oceanic influences: the mean annual temperature averages 9.2°C and total annual precipitation averages 820 mm. The oak-beech forest of Hesse is situated mainly on loamy or sandy soils moderately deep (24%) and on clay and loamy soils moderately deep (74%). An important windstorm occurred in 1999 with almost 12% of the forest destroyed, giving the opportunity of analyzing uprooted trees.

### 2.2 Tree selection

#### 2.2.1 Sample of trees for the aboveground biomass study

In order to represent the range of ages, trees were sampled in different plots of the forest over several years (Table 1). The trees first sampled (1996 and 1997) were previously analyzed for above and belowground biomass (Le Goff & Ottorini, 1998, 2001). The following samples allowed covering an age range from 8 to 172 years. In each sample, trees were selected so as to represent the different crown classes in the stand: dominant, co-dominant, intermediate and suppressed (Table 1).

**Table 1.** Description of the beech samples analyzed for biomass in Hesse forest (48°40’ N, 7°05’ E), from 1996 to 2003.

| Sample # | Forest plot # | Sample year | Stand age (1) | Sampled trees per crown class (2) |  |  |  | Total sampled trees (2) |
| --- | --- | --- | --- | --- | --- | --- | --- | --- |
|  |  |  |  | 1 | 2 | 3 | 4 |  |
| 1 | 217 | 1996 | 24 - 45 | 3 (3) | 3 (3) | 3 (3) | 2 (2) | 11 (11) |
| 2 | 217 | 1997 | 20 - 33 | 3 (1) | 3 (1) | 3 (1) | 3 (2) | 12 (5) |
| 3 | 222 | 2000 | 52 - 73 | 2 (2) | 2 (2) | 2 (2) | 0 (0) | 6 (6) |
| 4 | 215 | 2001 | 8 - 35 | 4 (3) | 5 (3) | 4 (4) | 4 (4) | 17 (14) |
| 5 | 214 | 2002 | 161 - 162 | 0 (0) | 1 (0) | 1 (1) | 0 (0) | 2 (1) |
| 6 | 220 | 2002 | 165 - 172 | 2 (1) | 0 (0) | 0 (0) | 0 (0) | 2 (1) |
| 7 | 220 | 2002 | 35 | 2 (0) | 0 (0) | 0 (0) | 0 (0) | 2 (2) |
| 8 | 215 | 2003 | 23 - 53 | 8 (0) | 1 (0) | 0 (0) | 0 (0) | 9 (0) |
| Total |  |  | 8 - 172 | 24 (10) | 15 (9) | 13 (11) | 9 (8) | 61 (40) |

(1) Stand age is given as the range of ages of the trees in each sample. (2) Sampled trees are sorted by crown classes: dominant (1), co-dominant (2), intermediate (3), suppressed (4) (Kraft classification, see Oliver & al, 1996); in parenthesis, the number of trees of the sub-sample analyzed for root biomass is given.

#### 2.2.2 Sub-sample of trees for the root biomass study

A sub-sample of trees was selected in almost each tree sample (#1 to #7) for root biomass analysis, so as the range of ages and the different social classes would be represented (Table 1). Then, a total of 40 trees – among the 61 trees sampled for the aboveground biomass analysis – were selected for root biomass analysis.

### 2.3 Pre-felling measurements

#### 2.3.1 Bole measurements

Several measurements were taken on sampled trees before they were felled. Breast height (1.3 m) was located on the stem and tree girth at breast height was measured. The upper end of the tree butt log was also located and the stem girth measured at this level.

#### 2.3.2 Crown measurements

The height to crown base^3^ was measured, either with a “*Vertex*” hypsometer (Pauwels, 2001) for tall trees or with a graduated telescopic pole for small trees. The horizontal crown projection was also established by measuring the distance and azimuth of the crown projection points^4^ from a fixed central point of the crown projection area; it allowed the calculation of the surface of the crown projection area of sampled trees.

### 2.4 Bole measurements

#### 2.4.1 In the field

After the measures on standing trees were performed, the trees were felled, while preventing them from falling brutally on the soil in the case of trees with limited dimensions. Tree heights for current and most recent years were measured with a tape whose graduation “1.3 m” was set at the breast height level marked on the trees; this allowed the calculation of current and most recent tree height increments. The crown base located on standing trees was marked on the stem and bole girth at this height was measured.

For each sample tree, sections of less than 1 m long were identified on the bole and arm forks, and the distances from stem (or arm fork) apex to the base of each stem (or arm fork) section were measured. After the stem and arm form sections identified were cut, the green weight of each section was measured, and a disc sample at the base of each section (10–15 cm thick) was taken off, identified and its green weight measured.

#### 2.4.2 In the laboratory

Ring radius measurements were performed on each stem disc collected in the field, in 4 perpendicular directions (8 directions at 45° for irregular discs) and for the last 6 years, allowing the calculation of the radial increment of the last 5 years for each disc. A ring count on the disk taken at stump level was also performed to obtain tree age.

The green weights of a sub-sample of the stem discs were measured, with and without bark. Moreover, the dry weights of the sampled and sub-sampled stem disks were obtained by leaving the wood samples in a drying oven at 105°C until the weight was stabilized; for the sub-samples, wood and bark were weighted separately.

### 2.5 Branch measurements

#### 2.5.1 In the field

For each sample tree, branches were identified, except the very small ones that were grouped. As for the bole, sections of length less than 1m were delimited and identified on each branch of length larger than 1m. The green weight of each branch, or branch section, was measured, and a disc sample at the base of each branch section –10 to 20 cm long – was taken off, identified and its green weight measured.

Dead branches, epicormics and beechnuts were harvested and grouped separately. For each branch and group of branches and epicormics, the twigs and their leaves were harvested and their green weight measured as a whole, and a sample taken off and put in a closed plastic bag to prevent desiccation.

#### 2.5.2 In the laboratory

The following measures were obtained:

-green weights of twigs and leaves measured separately for each sample collected in the field,
-area of leaves for each sample by using a scanner and *ImageJ* (http://rsb.info.nih.gov/ij/) software which gives individual leaf area values and total leaf sample area,
-green weights of wood and bark for a sub-sample of branch discs,
-ring widths in 4 perpendicular directions for the last 6 years on each branch disc,
-dry weights of branch discs (separately for wood and bark for the sub-samples), of twigs and leaves samples (separately) and of beechnuts, after drying at 105°C for 48 hours.

### 2.6 Adaptations to stem and branch measurements

#### 2.6.1 Large trees

Several adaptations to the protocol of stem and branch measurements were made in the case of large trees (samples #3 & #5: see Table 1).

##### Stem measurements

-the lower part of the bole (0–1.30 m) was divided in 3 parts (0–0.30 m; 0.30–0.80 m; 0.80–1.30 m) to take account of the butt swell,
-the basal and top girths of bole and forks sections were measured to estimate their volumes, and a disc sample was taken at the base of each section allowing to convert volume to dry weight,
-the bole and fork ends were isolated and treated as “small trees” with the same processes and measurements,
-in the case of bole or fork ends breakage at the time of tree felling, a special measurement process was applied as for broken branches (see after).

##### Branch measurements

-branches were inventoried and their basal diameter or girth measured (in two perpendicular directions for diameter), as well as their length and height of insertion on tree bole,
- 3 branches per each quarter of the crown were sampled and treated as “small trees”
- in the case of branch breakage at the time of tree felling, an intact section of comparable diameter – basal diameter of the intact section comparable to the diameter at the end of the broken branch – on the same or another branch was selected so as to estimate the missing branch, twigs and leaves biomasses of the broken branch.

#### 2.6.2 Sample # 8 (see Table 1)

This additional sample of trees was taken in the same plot as sample # 4 in order to extend the range of already observed tree diameters with trees of larger diameter and to better fit the diameter distribution of the stand.

The same measurement process as for large trees was applied:

-stem sections were defined on the bole so as to evaluate the volume and the volume increment of the stem and a disk sample was taken on each stem section to estimate stem biomass and biomass increment,
-a diameter inventory of branches was performed and a sample of branches (3 per each half part of the crown in this case) was selected to establish biomass equations.

Additional measurements^5^ were also taken on the stem and branches, in order to evaluate the biomass characteristics for year 2001 (as for sample # 4), in addition to those of year of tree felling (2003):

-height reached by the stem and fork arms in 2001,
-diameter inventory of branches distinguishing the branches formed after 2001 (situated on stem growth units of years 2001 and later) that will not be considered in the calculation of tree branch biomass and biomass increment of year 2001,
-on sampled branches:
  + under bark basal diameter in 2001, allowing to establish a relation with basal diameter in 2003 to be applied to the inventory of branches for year 2001,
  + taking of disc samples (1 or 2 per branch depending on branch size) to evaluate the biomass increment of sampled branches in 2003 and 2001 from branch biomass and relative branch section area increment obtained from disc samples.

### 2.7 Root measurements

#### 2.7.1 Root system extraction

The root systems were excavated using a mechanical shovel and an air knife (more specifically for uprooted trees) so as to minimize the loss or breakage of roots. Then, the root systems were washed to remove remaining soil particles and finally dried in open air. In case of root breakage, the root parts left in the soil were extracted so as to reconstruct the root system. In case of very large root systems (as for trees of sample # 5), a special apparatus was constructed to lift up the already extracted roots, allowing the extraction of the remaining part of the root system still in the soil.

#### 2.7.2 Measurement processing

The measurement process used for tree samples #1 & # 2 (see Le Goff & Ottorini, 2001) was applied to the trees sampled later:

-roots were sorted into 3 size classes depending on the cross-sectional diameter (*d*) of the roots : coarse roots (*d* ≥ 5mm), small roots (2 ≤ *d* < 5 cm), fine roots (*d* < 2 mm).
-on coarse and small roots, root samples were taken (about 10 cm long and of regular shape) to estimate the current annual volume and biomass increments of the root system; the number of increment samples per tree varied from 2 to more than 30, depending on tree dimensions which vary with age and social status.
-the length, the diameter at both ends along two perpendicular directions and the annual increments every 45° of each root sample were measured; after synchronization of the annual increments, the mean annual radial increment and the annual volume increment of each root sample were calculated.
-the diameters of broken root ends were measured outside bark at the point of breakage in two perpendicular directions including the major axis.
-unbroken root ends were sampled for estimating the missing biomass of broken roots : 2 perpendicular diameters at various cut ends of unbroken root ends were measured outside bark so as to establish a regression of root weight on root end diameter and apply it to broken root ends measured in diameter (Le Goff & Ottorini, 2001).
-the root systems were oven-dried to a constant weight at 105°C, and the dry weight of each root category – coarse, small and fine – was recorded separately for each root system and for each unbroken root end. The same drying process was used to obtain the dry weights of the root samples.

### 2.8 Data processing

#### 2.8.1 Stem

##### 2.8.1.1 Volume and volume increment

When stem scaling was performed, the stem volume was obtained as the sum of the volumes of the different sections making the stem. Each section was considered as a truncated cone, except the last one on stem ends that was considered as a cone. In this case, the current volume increment of stem sections was calculated as the product of the stem volume sections by the relative current area increment of the increment samples taken on each stem section.

When stem scaling was not performed (sample # 1), the stem volume was obtained by converting dry weights of stem sections into volume using the specific gravity (ratio of dry weight to volume) of the sampled stem discs. In this case, bole volume increment was derived from bole biomass increment, using a relation established on the whole set of trees subjected to stem scaling.

##### 2.8.1.2 Biomass and biomass increment

Stem biomass was calculated as the sum of the dry weights of the bole and of the fork arms sections, the dry weight of each section being generally evaluated as the product of the green weight of the section by the ratio of dry weight to green weight of sample disks taken in each stem section. When samples were not available for each stem section, a relation was established on sample disks relating wood specific gravity to the height of sample disks and the biomass of stem sections without samples was then estimated as the product of stem section volume by the specific gravity estimated from the mean height of stem section.

The weighting separately of wood and bark for a sub-sample of stem disks allowed to estimate the dry weight of wood and bark of the stem sections and of the whole stem, using a relation describing the variation of the ratio of bark (or wood) to total dry weight along the stem for the sub-sample of stem discs.

In case green weights were not measured in the field, the dry weight of each stem section was calculated as the product of the stem section volume by the specific gravity of the disc sample taken in the stem section or estimated as above in case of missing sample.

The current biomass increment of stem sections was calculated, as for current volume increment, as the product of stem sections biomass by the relative current area increment of the increment samples taken in each stem section. When increment samples were not available for each stem section, a relation was established relating the relative current area increment of available increment samples to their height (or relative height) in the tree.

##### 2.8.1.3 Stem area

When stem scaling was performed, the stem area of trees was obtained as the sum of the areas of the different sections making the stem as for bole volume.

When stem scaling was not performed (as for sample #1), the stem area of trees was derived from bole biomass, using a relation established on the whole set of trees subjected to stem scaling.

#### 2.8.2 Branches

##### 2.8.2.1 Basal diameter

The basal diameter of branches was calculated as the geometric mean of the 2 diameters measured at right angles at the base of branches.

##### 2.8.2.2 Biomass and biomass increment

In case of a complete inventory of branches on sampled trees, the dry weight of each branch was calculated as the sum of the dry weights of the branch sections estimated as the product of sections green weights by the mean ratio of dry weight to green weight calculated for the whole set of branch samples. In case of branch sampling, relations were established at tree level linking branch biomass to branch basal diameter and these relations were used to estimate the biomass of non-sampled branches.

The total dry weight of branches per tree was then calculated as the sum of individual branch dry weights, of the dry weight of grouped small branches (obtained as for sampled branches), and of the dry weight of stem ends branches and of epicormics for concerned trees (the dry weight of grouped epicormics being obtained as for sampled branches).

The weighting separately of wood and bark for a sub-sample of branch disks allowed estimating a mean branch wood biomass ratio for each tree. This ratio was used to estimate the wood biomass of each branch from its total biomass and the bark biomass was obtained as the difference between total and wood biomasses.

The current annual biomass increment of branch sections was calculated, as for stem biomass increment, as the product of branch sections biomass by the relative current area increment of the increment samples taken in the corresponding branch sections. For trees with sampled branches, relations were established between branch biomass increment and either branch basal diameter (tree sample # 3) or foliage biomass (tree sample # 5); these relations were used to estimate the biomass increment of non-sampled branches. The current biomass increments of grouped branches (branches on stem ends and groups of small branches) were calculated as the product of branch groups biomasses by the relative current area increment of the increment samples taken in each group.

#### 2.8.3 Leaves

##### 2.8.3.1 Dry weight

###### branch leaf dry weight

In case of a total branch inventory of leaves with their supporting twigs (tree samples #1 & #2), branch leaf dry weights were obtained by multiplying the “leaf+twigs” green weight by the ratio of sample leaf dry weight to sample “leaf+twigs” green weight. In this case, leaf dry weight was calculated separately for shade and sun leaves.

In case of leaves and twigs collected together for all branches, except for a sample of branches^6^ for which “leaf+twigs” samples were taken (tree samples # 4, # 7) allowing to calculate their leaf dry weight as before, the leaf dry weight of grouped branches was obtained by multiplying their total “leaf+twigs” green weight by the mean ratio of leaf dry weight to green weight of “leaf+twigs” samples.

In case of “leaf+twigs” collected only for a sample of branches (tree samples # 3 & # 5), after estimating the leaf dry weight of sampled branches from a “leaf+twigs” sample as before, relations were established at tree level linking branch leaf dry weight to basal branch diameter (quadratic or power model) or to dry weight of branches (power model). These relations were then used to estimate the leaf dry weight of non sampled branches.

###### stem ends leaf dry weight

For each stem end defined (tree samples # 3 & #5), the total leaf dry weight was calculated from total “leaf+twigs” green weight, leaf dry weight and “leaf+twigs” green weight of a “leaf+twigs” sample, as for sampled branches, and the leaf dry weights of all stem ends were summed.

###### epicormics leaf dry weight

For each set of epicormics recognized, the total leaf dry weight was calculated from total “leaf+twigs” green weight, leaf dry weight and “leaf+twigs” green weight of a “leaf+twigs” epicormics sample.

The total tree leaf dry weight was then calculated as the sum of the leaf dry weight of branches, leaf dry weights of stem ends (if any) and leaf dry weights of epicormics (if any).

##### 2.8.3.2 Leaf area

###### branch leaf area

In case of “leaf+twigs” samples taken on each inventoried branch (tree samples # 1 & # 2), total leaf branch area was obtained by multiplying the leaf sample area by the ratio of branch leaf dry weight (measured or estimated) to leaf sample dry weight.

In case of leaf+twigs collected together for all branches, except for a sample of branches for which leaf+twigs samples were taken (tree samples # 4, # 7) allowing to calculate their leaf area as before, the leaf area of grouped branches was obtained by multiplying the total calculated leaf dry weight by the mean ratio of leaf area to leaf dry weight (SLA) of the samples taken in each crown stratum.

In case of “leaf+twigs” collected only for a sample of branches (tree samples # 3 & #5), after estimating the leaf area of sampled branches as before, relations were established at tree level (tree sample # 5) or for the whole sample of trees (tree sample # 3), linking directly total branch leaf area to basal branch diameter. In case of sample #3, the equation established of form ln(y) = a + ln(x) was inversed and corrected for bias; in case of sample # 5, the equation was a quadratic polynomial satisfying that leaf area is null when basal branch diameter is 0.

###### stem ends leaf area

For each stem end defined (tree samples # 3 & # 5), total leaf area was obtained by multiplying the leaf sample area by the ratio of stem end leaf dry weight to leaf sample dry weight.

###### epicormics leaf area

For each set of epicormics recognized, total leaf area was obtained by multiplying the leaf sample area by the ratio of epicormics leaf dry weight to leaf sample dry weight.

The total tree leaf area was obtained as the sum of leaf area of branches, leaf area of stem ends (if any) and leaf area of epicormics (if any)

#### 2.8.4 Twigs

The same process as for leaves was followed to obtain the dry weight of branch twigs, replacing leaf weight by twigs weight in the calculations performed or in the relations to be established to estimate the total dry weight of twigs from basal branch diameter.

#### 2.8.5 Roots

##### 2.8.5.1 Biomass

Biomass equations were established for broken roots on a sample of intact root ends (see Le Goff & Ottorini, 2001) in order to estimate total missing root biomass, with distinction made between horizontal and vertical roots for larger trees (samples # 3 and # 5), and in order to estimate the missing root biomass for each root fraction of each category of roots.

The total tree root biomass was obtained as the sum of the dry weights calculated for each root category, for measured roots, sampled root ends, root increment samples and missing root ends.

The missing root biomass represented between 0 and 10% of the total root biomass of trees (sample # 4), between 2 and 7% (sample # 5) and between 5 and 20% (sample # 3).

##### 2.8.5.2 Biomass increment

The calculation of the current annual biomass increment of the root systems was based on the current annual relative volume increments of the root increment samples taken, with the following steps:

-calculation of inside bark current annual relative volume increments (dv/v)_n_ of the root increment samples, where v is the volume of the increment sample for current year n. The root increment samples were considered as truncated cones of height “l”, length of the increment sample, and mean inside bark diameters D_q_ and d_q_ for the sides of larger and lower diameters respectively^7^. The current annual relative volume increments (dv/v)_n_ were calculated as Log v_n_ – Log v_n−1_, where v_n_ and v_n−1_ are the volumes of the increment samples for years n and n−1 respectively.
-calculation of the median (k) of the current annual relative volume increments (dv/v)_n_ of the root increment samples of trees for each crown class (see Le Goff & Ottorini, 2001), the current annual relative volume increments being independent of the cross sectional area of increment samples. Depending on tree sample, k appeared dependent or not on crown class (Table 2). **Table 2.**
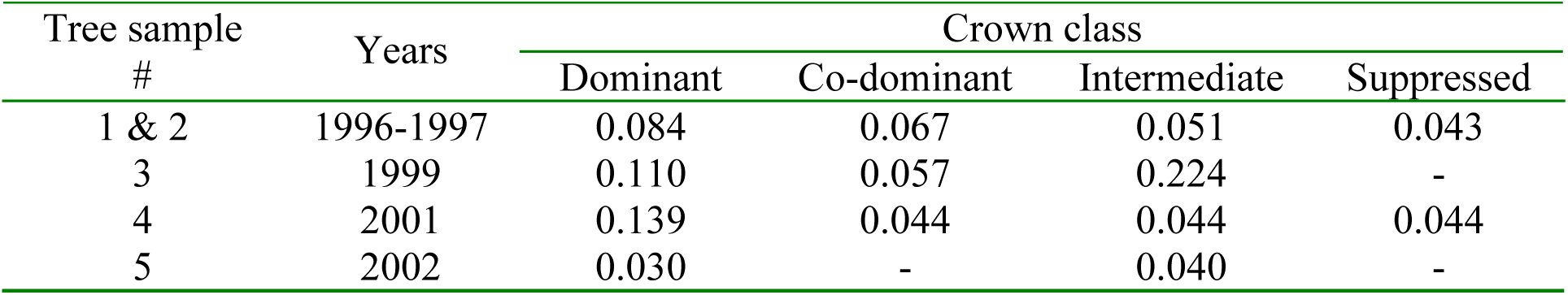
Median values (k) of the current annual relative volume increments of the root increment samples of sampled trees per crown class.

The median values (k) of current relative annual volume increments were used as estimates of the relative annual volume increments of the whole root systems of trees (see Le Goff & Ottorini, 2001) in each crown class. The current annual biomass increments of large and small roots were then obtained by multiplying their biomass by the appropriate “k” value, considering that the wood density of all parts of the root system was constant (see Le Goff & Ottorini, 2001).

In this study, the fine root turnover was not considered; it was estimated around 0.6 t ha^−1^ an^−1^ at stand scale for the experimental stand “Hesse-1” at the age of 20 with a stand density of about 3500 stems ha^−1^, which corresponds to about 0.17 kg an^−1^ at tree scale (Le Goff & Ottorini, 2001).

### 2.9 Biomass analysis

#### 2.9.1 Biomass models

##### selection of a generic model for biomass

The following model^8^ already retained by Genet & al (2011a), and recommended recently by McElligott & al. (2013) for its good extrapolation properties, was selected to represent the variations of the biomass (*W*) of the different tree compartments (taken alone or grouped) with measured tree attributes, that is breast height diameter (*D*) and total height (*Ht*):

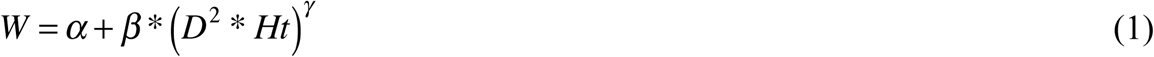

In this equation, the parameter α is generally non significant (Genet & al, 2011b). Then, the model described by Eq. 1 could be reduced to the following equation:

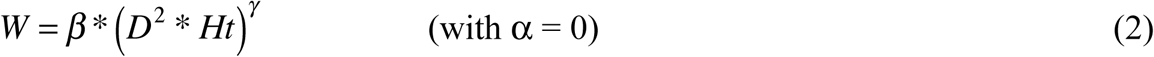

Moreover, in Eq. 2, the parameters β and *γ* may vary with other tree attributes such as age or competitive status, or depend on tree stand belonging.

##### model fitting

Two procedures were used for fitting Eq. 2 and then compared. First, the fitting of Eq. 2 was done after a “both sides” logarithmic transformation allowing the use of linear regression; second, Eq. 2 was fitted directly using a non-linear regression procedure.

In the first procedure, the log-transformation of both sides of Eq. 2 gives:

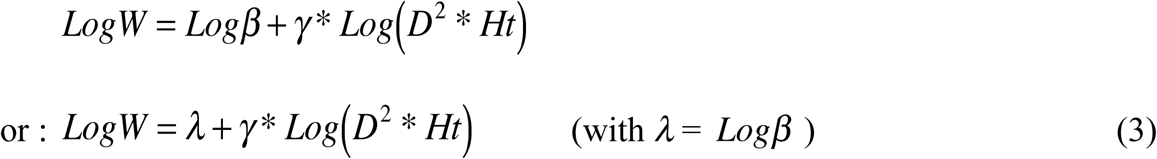

Eq. 3 was fitted using the software *Data Desk 6.3* (Velleman, 2011) on a Mac OS 10 system, to total above ground biomass for each age class, after trees were sorted into 4 age classes (Table 3). An analysis of the variation of each parameter (*λ*, *γ*) with age (*Age*) and relative crown length (*RCL*)^9^ was then performed, *λ* and *γ* being linearly related to *RCL* and to the inverse of *Age.*

**Table 3.** Distribution of tree samples into 4 age classes, with corresponding mean age.

| Age class | Tree samples<br># | Mean age<br>(years) |
| --- | --- | --- |
| 1 | 4 | 21.7 |
| 2 | 1, 2, 7, 8 | 31.6 |
| 3 | 3 | 58.8 |
| 4 | 5, 6 | 165.0 |

<sup>8</sup> In this paper, as recommended by Sileshi (2014), it will not be referred to “allometric models” to design the biomass equations used, as they don’t follow the typical power-law function.
<sup>9</sup> The relative crown length (*RCL*) of a tree was defined as the ratio of crown length (distance from crown base to stem apex) to total tree height (*Ht*).

The following equation, derived from Eq. 3 and including *Age* and *RCL* variables, was then fitted to biomass data:

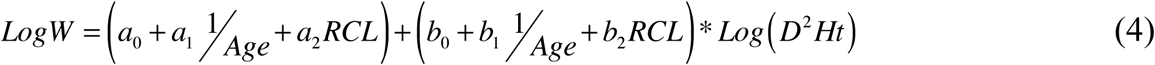

Equation (4) was fitted with *Datadesk 6.3* to each above (stem and branches) and belowground (coarse, small and fine roots) tree compartment and to total above and total belowground biomasses – with possible adaptations to Eq. 4 – and the residuals of this equation were examined to detect any bias or remaining tree or stand effect.

In the second followed procedure, Equations (1) or (2) were fitted as non-linear models, using the *R* project for statistical computing (2009), with parameters dependent more specifically on tree age (Genet & al, 2011). The fitted equation was the following:

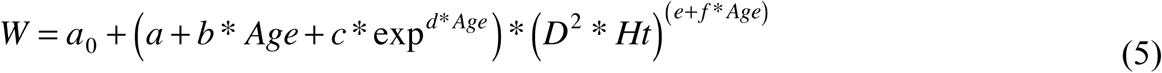

Equation (5) was fitted to each above (stem and branches) and belowground (coarse, small and fine roots) tree compartment and for total above and total belowground biomasses with the statistical package *nls* of *R* (2009), and the residuals were examined as for the first procedure.

##### simultaneous model fitting

A multivariate procedure was applied in order to take account of the statistical dependence among the biomass equations of the different tree compartments. In this way, multivariate models are able to ensure a better additivity of tree compartments, compared to the independently estimated equations, without the need to address a constraint of additivity in the models (Repola, 2009).

The multivariate models were fitted separately for above and belowground tree compartments as the number of observations diverged due to the sub-sampling procedure applied for belowground biomass.

The multivariate procedure consisted in fitting the biomass equations by using dummy variables (Zeng & al, 2011) to render the parameters of the equations specific of each tree compartment (Repola, 2008, 2009). In case of non-linear fitting, the estimated values of the parameters obtained by separate fittings were used as starting values in the simultaneous fitting process.

##### random effects

The residuals of the simultaneous fitting of biomass equations were examined with particular attention for a possible remaining “stand” or “tree status” (crown class) effect, in relation with the sampling scheme. In such case, a mixed model procedure, using the *lme* or *nlme* package of *R*, was applied to take account of the above random effects in the biomass equations and an analysis of the residuals was performed to verify that no such remaining effect was detectable.

##### selection of model parameters and quality of model predictions

Uncertainty around fixed model parameters was evaluated by considering the percent relative standard error (*PRSE*) defined as
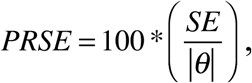
where *θ* is the estimated value of a given parameter and *SE* its corresponding standard error: point estimates of *θ* are generally considered unreliable if PRSE > 25% (Sileshi, 2014).

To evaluate the quality of the fitted biomass models, the observed values were plotted and regressed against the predicted values (Pineiro et al., 2008; Sileshi, 2014), and the regression lines were compared graphically with the 1:1 lines (for a good model, the coefficients of the regression lines would be close to 0 for the constant and 1 for the slope).

#### 2.9.2 Biomass increment models

Two alternative models were proposed by Hofmeyer & al. (2010) in order to represent the relationship between annual stem volume increment and tree leaf area. The following exponential model (Eq. 6) proved better adapted than the allometric model as it was in accordance with the non-monotonically variation of stem growth efficiency^10^ with tree leaf area, as observed in our case. Then, this model was retained here to explore the variations of the biomass increment (Δ*W*) of the different tree compartments with tree leaf area (*LA*).

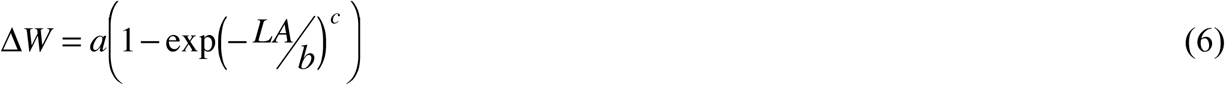

The possible dependence of Δ*W* on other tree characteristics measured was explored by examining the residuals of Eq. 6 (for additive effects) and the ratios of the observed and estimated biomass increments (for multiplicative effects).

As for biomass equations, multivariate non-linear models were fitted separately for above and below ground tree compartments, and separate fittings were performed for each compartment to obtain starting values of the model parameters when fitting the multivariate models. Mixed models were fitted with *nlme* (*R*) to test possible random effects due to “forest plot”, “year of sampling” or “tree social status”. Weightings (specific of each compartment) and correlations were also considered when fitting simultaneously the biomass increment models for above and below ground tree compartments: in this scope, an indicator variable was introduced in the data set to identify each tree compartment (Repola, pers. comm.).

As for biomass equations also, percent relative standard errors (*PRSE*) were considered to judge of the uncertainty around fixed model parameters, and observed against predicted biomass increment values were plotted and regressed to assess the quality of the models.

## 3. Results

### 3.1 Tree sample characteristics

#### 3.1.1 Aboveground tree sample (Table 4)

The main characteristics of the tree sample which are listed in Table 4 let appear the wide range of observed values for the aerial part of trees, due mainly to the large range of the tree ages and of the competitive status of trees (crown ratio – the ratio of crown length to total tree height (RCL) – varying between 0.2 and 0.9); this is particularly the case for biomass values which vary in the proportion of more than 1 to 10000.

**Table 4.** Descriptive attributes of the 61 destructively sampled beech trees in Hesse forest (NE France) analyzed for aboveground biomass and representing different age and social status classes.

| Attributes | Nb. of trees<br>(1) | Mean (SE) | Range |
| --- | --- | --- | --- |
| Age (years) | 61 | 40.3 (35.5) | 8 - 172 |
| DBH (cm) | 61 | 10.5 (11.4) | 1.1 – 60.5 |
| Height (m) | 61 | 12.2 (8.3) | 2.2 – 39.3 |
| Crown length (m) | 61 | 5.9 (4.4) | 1.0 – 24.2 |
| Crown ratio | 61 | 0.50 (0.16) | 0.22 – 0.89 |
| Crown projection area (m<sup>2</sup>) | 39 | 9.77 (20) | 0.85 – 101,53 |
| Stem area (m<sup>2</sup>) | 61 | 5.09 (11.64) | 0.08 – 66.56 |
| Stem volume (m<sup>3</sup>) | 61 | 0.2802 (0.9373) | 0.0002 – 5.8775 |
| Leaf area (m<sup>2</sup>) | 59 | 46.74 (102.03) | 1.11 – 709.63 |
| LAI (m<sup>2</sup> m<sup>-2</sup>) | 39 | 3.62 (1.59) | 1.30 – 7.98 |
| Leaf mass (kg) | 59 | 2.057 (4.493) | 0.008 – 31.523 |
| Branch mass (kg) | 61 | 26.620 (79.096) | 0.026 – 493.856 |
| Stem mass (kg) | 61 | 162.434 (545.336) | 0.126 – 3442.037 |
| Stem volume increment (m<sup>3</sup> an<sup>-1</sup>) | 61 | 0.011 (0.022) | 0.000 – 0.112 |
| Stem mass increment (kg an<sup>-1</sup>) | 61 | 6.027 (12.179) | 0.003 – 61.197 |
| Branch mass increment (kg an<sup>-1</sup>) | 61 | 1.759 (3.916) | 0.001 – 21.916 |

(1) Crown projection area could not be measured on all trees and then this variable, as well as leaf area index (LAI), were only available for 39 trees. Leaf area and biomass could not be obtained for 2 trees and were then available for 59 trees only.

#### 3.1.2 Above and belowground tree sample (Table 5)

The belowground attributes, together with the aboveground ones, were only measured on a sub-sample of 40 trees. However, the range of observed values for above and belowground attributes of the sub-sample remains large, as the range of tree ages is the same as for the complete sample (Table 3). Thus, the total root biomass of sampled trees varies in the proportion of 1 to 5000, while the total aboveground biomass still vary in the proportion of 1 to 10000.

**Table 5.**
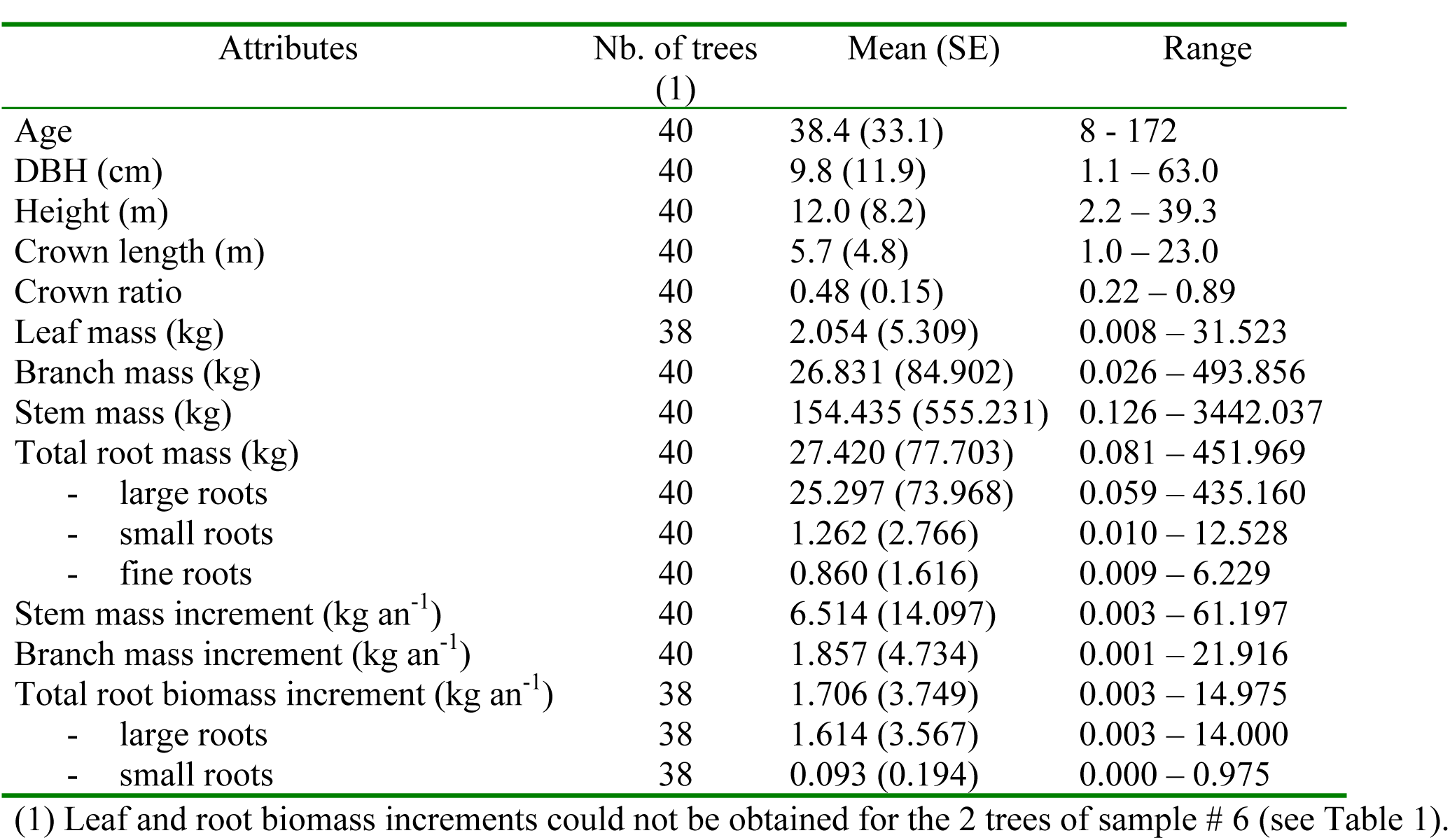
Descriptive attributes of the sub-sample of 40 destructively sampled beech trees in Hesse forest (NE France) analyzed for root and aerial biomass and representing different age and social status classes.

| Attributes | Nb. of trees<br>(1) | Mean (SE) | Range |
| --- | --- | --- | --- |
| Age | 40 | 38.4 (33.1) | 8 - 172 |
| DBH (cm) | 40 | 9.8 (11.9) | 1.1 – 63.0 |
| Height (m) | 40 | 12.0 (8.2) | 2.2 – 39.3 |
| Crown length (m) | 40 | 5.7 (4.8) | 1.0 – 23.0 |
| Crown ratio | 40 | 0.48 (0.15) | 0.22 – 0.89 |
| Leaf mass (kg) | 38 | 2.054 (5.309) | 0.008 – 31.523 |
| Branch mass (kg) | 40 | 26.831 (84.902) | 0.026 – 493.856 |
| Stem mass (kg) | 40 | 154.435 (555.231) | 0.126 – 3442.037 |
| Total root mass (kg) | 40 | 27.420 (77.703) | 0.081 – 451.969 |
| - large roots | 40 | 25.297 (73.968) | 0.059 – 435.160 |
| - small roots | 40 | 1.262 (2.766) | 0.010 – 12.528 |
| - fine roots | 40 | 0.860 (1.616) | 0.009 – 6.229 |
| Stem mass increment (kg an<sup>-1</sup>) | 40 | 6.514 (14.097) | 0.003 – 61.197 |
| Branch mass increment (kg an<sup>-1</sup>) | 40 | 1.857 (4.734) | 0.001 – 21.916 |
| Total root biomass increment (kg an<sup>-1</sup>) | 38 | 1.706 (3.749) | 0.003 – 14.975 |
| - large roots | 38 | 1.614 (3.567) | 0.003 – 14.000 |
| - small roots | 38 | 0.093 (0.194) | 0.000 – 0.975 |

(1) Leaf and root biomass increments could not be obtained for the 2 trees of sample # 6 (see Table 1).

### 3.2 Above and belowground biomass

#### 3.2.1 Biomass distribution (in relation to age and social status of trees)

The variations of the tree biomass distribution among above and belowground tree compartments were observed by splitting sample trees in different social and age classes. Social classes refer to the classical Kraft classification (Oliver and Larson, 1996) which differenciates dominants, co-dominants, intermediate and suppressed trees. The sample trees were sorted into 4 age classes (numbered 1, 2, 3, 4) with mean ages 22, 32, 59 and 165 respectively.

##### aboveground biomass

The distribution of the aboveground biomass among tree compartments showed the same pattern with age, independently of tree social class (Fig. 1): the proportion of aboveground tree biomass corresponding to stem wood – between 60 and 80% – increased with age, while, at the same time, it decreased for leaves and seemed to peak at intermediate ages for branches.

**Figure 1.**
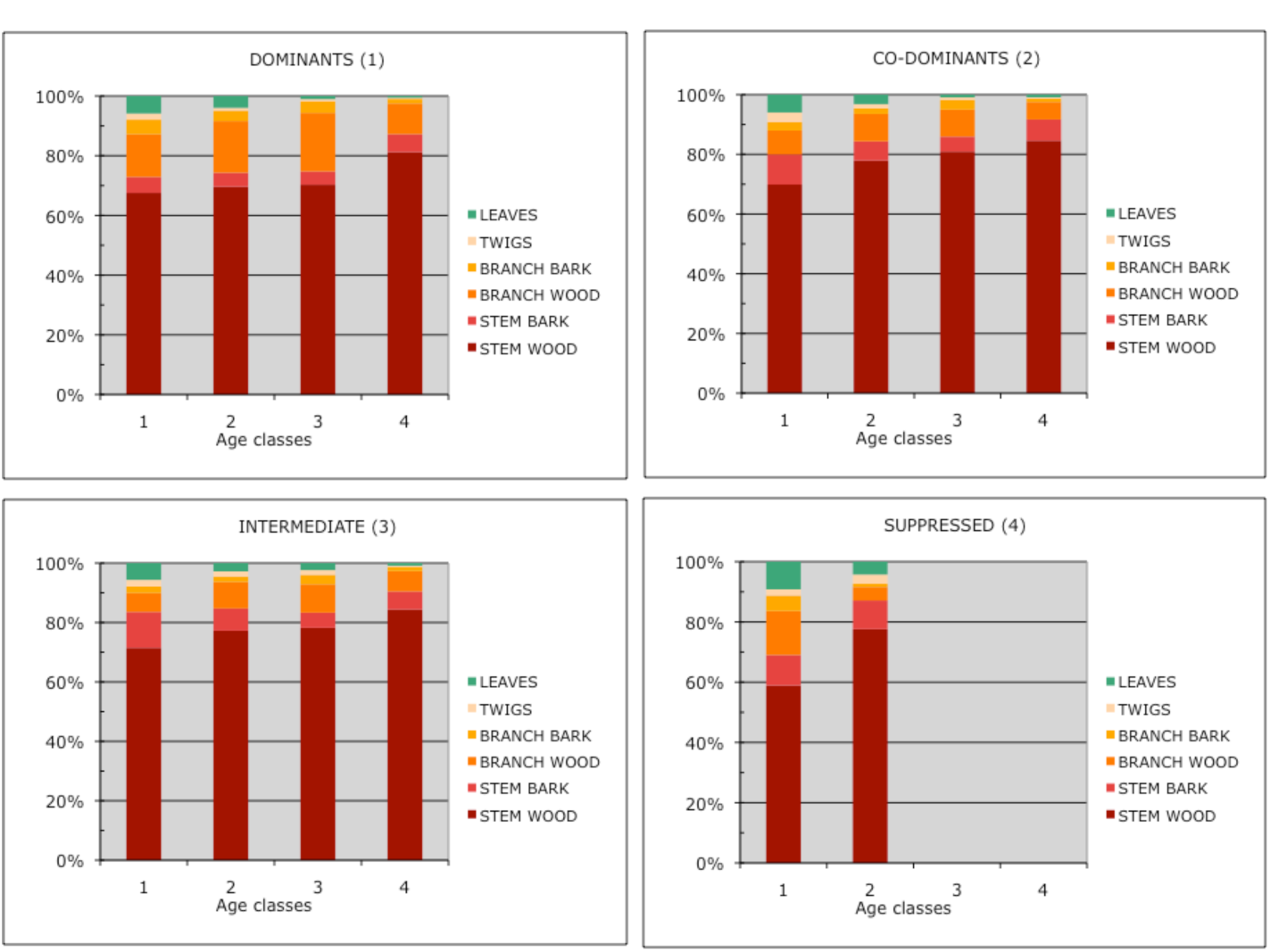
Distribution (in %), by age and social classes, of the aboveground tree biomass among the following tree compartments: leaves, twigs, branches (bark and wood) and stem (bark and wood).

##### above and belowground biomass

As shown by Fig. 2, the proportion of tree biomass present in roots tended to decrease with age, from nearly 20% to 10% or less. The biomass of the root system appeared comparable to that of the branch compartment, but seemed generally slightly larger for a given age class, except for dominant trees in the two highest age classes.

**Figure 2.**
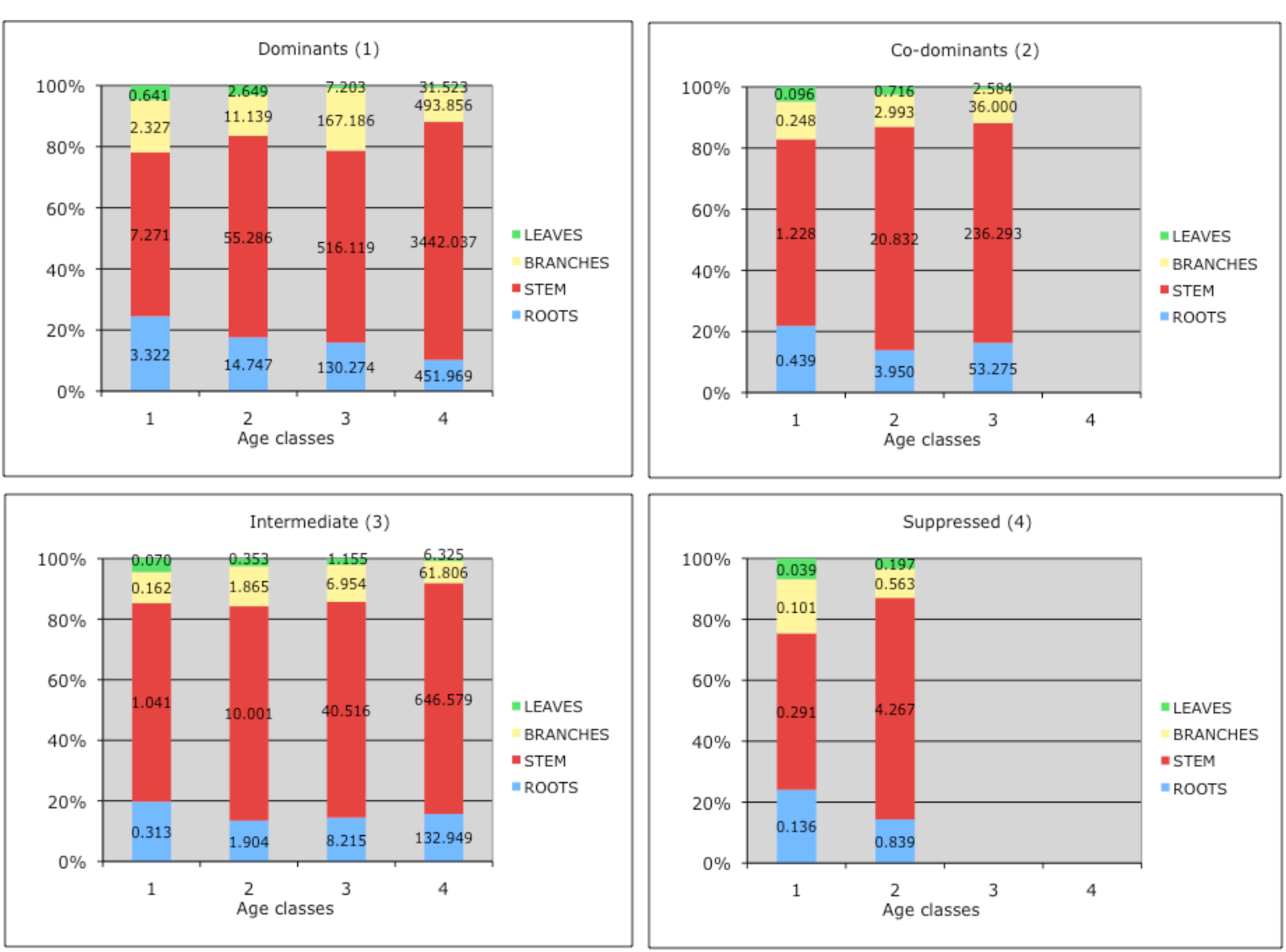
Distribution (in % and kg), by age and social classes, of the above and belowground tree biomasses among the main tree compartments: leaves, branches (including twigs), stem and roots (bark added to wood for stem and branches).

#### 3.2.2 Biomass equations

##### aboveground

Biomass equations were established for the following tree compartments: stem (wood + bark), branches (wood + bark) + twigs called all together “branches”, total aboveground (stem + “branches”)

The non-linear fitting of Equation (5) gave better results than the linear fitting of the log-transformed Equation (3) with which biased estimations were observed (even after correction for inverse log transformation). The simultaneous estimation of the parameters retained after a separate fitting of the equations for each tree compartment, produced the following results (Table 4):

**Table 4.**
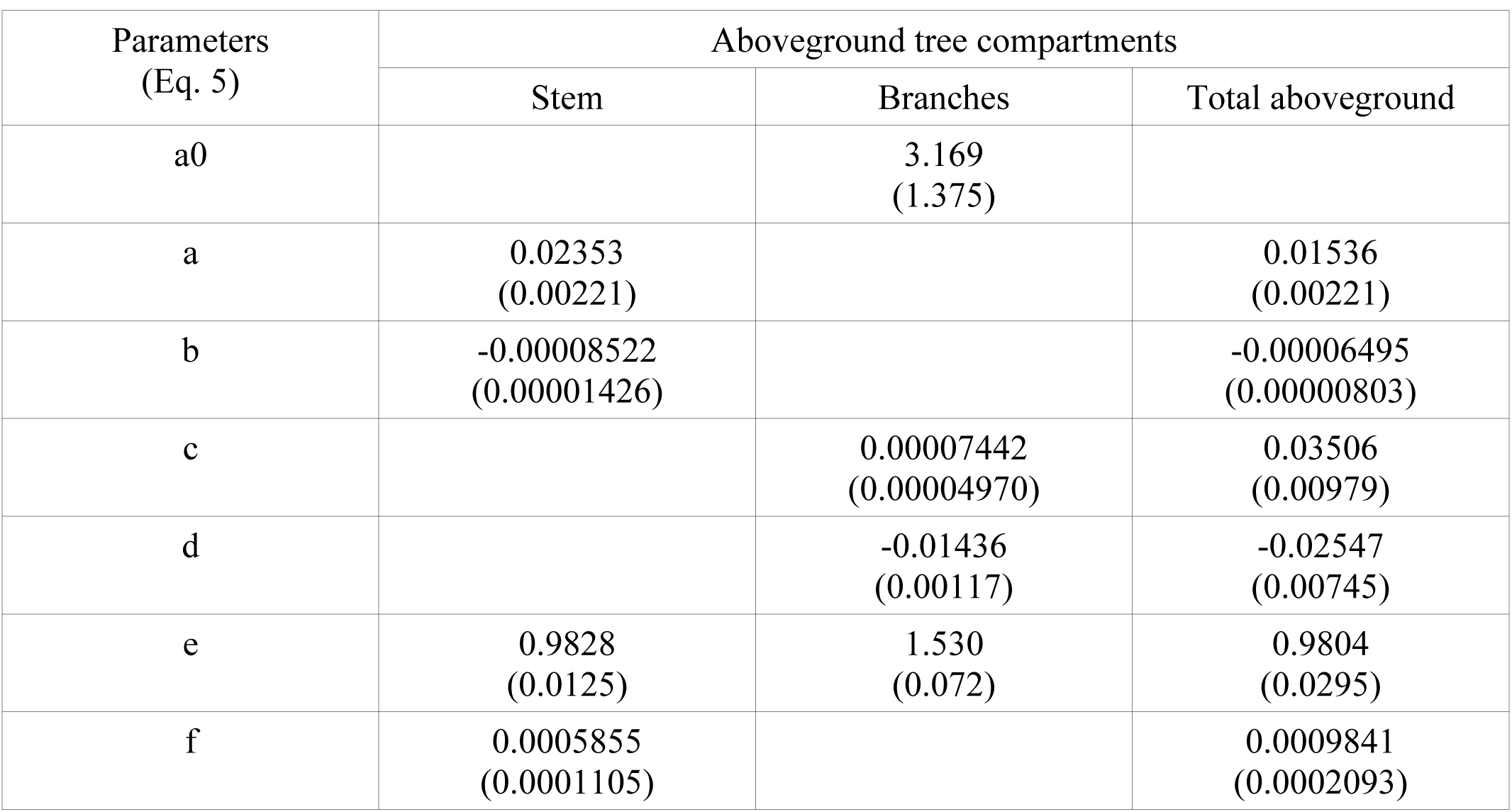
Estimated values of the parameters of Equation 5, with their standard errors (in parenthesis), for each aboveground tree compartment (stem and branches) and for the total aboveground biomass. (*all parameters are significant at the 0.05 level or more, except the parameter “c” for branches which was nevertheless retained in Eq. 5 as it was significant when fitting separately the branch biomass equation*)

| Parameters<br>(Eq. 5) | Aboveground tree compartments |  |  |
| --- | --- | --- | --- |
|  | Stem | Branches | Total aboveground |
| a0 |  | 3.169<br>(1.375) |  |
| a | 0.02353<br>(0.00221) |  | 0.01536<br>(0.00221) |
| b | -0.00008522<br>(0.00001426) |  | -0.00006495<br>(0.00000803) |
| c |  | 0.00007442<br>(0.00004970) | 0.03506<br>(0.00979) |
| d |  | -0.01436<br>(0.00117) | -0.02547<br>(0.00745) |
| e | 0.9828<br>(0.0125) | 1.530<br>(0.072) | 0.9804<br>(0.0295) |
| f | 0.0005855<br>(0.0001105) |  | 0.0009841<br>(0.0002093) |

The observation of the residuals of Eq. 5 did not reveal any bias in the biomass estimation of each compartment, nor any relation with tree variables not already entered in the equation. Moreover, the plotting of residuals for trees belonging to the different forest plots or social classes sampled did not reveal any marked dependence. This was confirmed by the limited effects observed when adding random effects in Eq. 5 – fitted in this case with *nlme* (R): among all possible random effects associated with each parameter of Eq. 5, a significant one associated with “forest plot” was only observable for the parameter “*f*” of stem biomass equation, and it was then decided to ignore random effects when fitting Eq. 5.

The estimated biomass of the different aerial tree compartments compared favorably with the observed biomass values (Figures 3A, 3B, 3C). Moreover, the sum of the estimated stem and branch biomasses compared relatively well with the total aboveground biomass estimated as a whole (Figure 3D)

**Figure 3.**
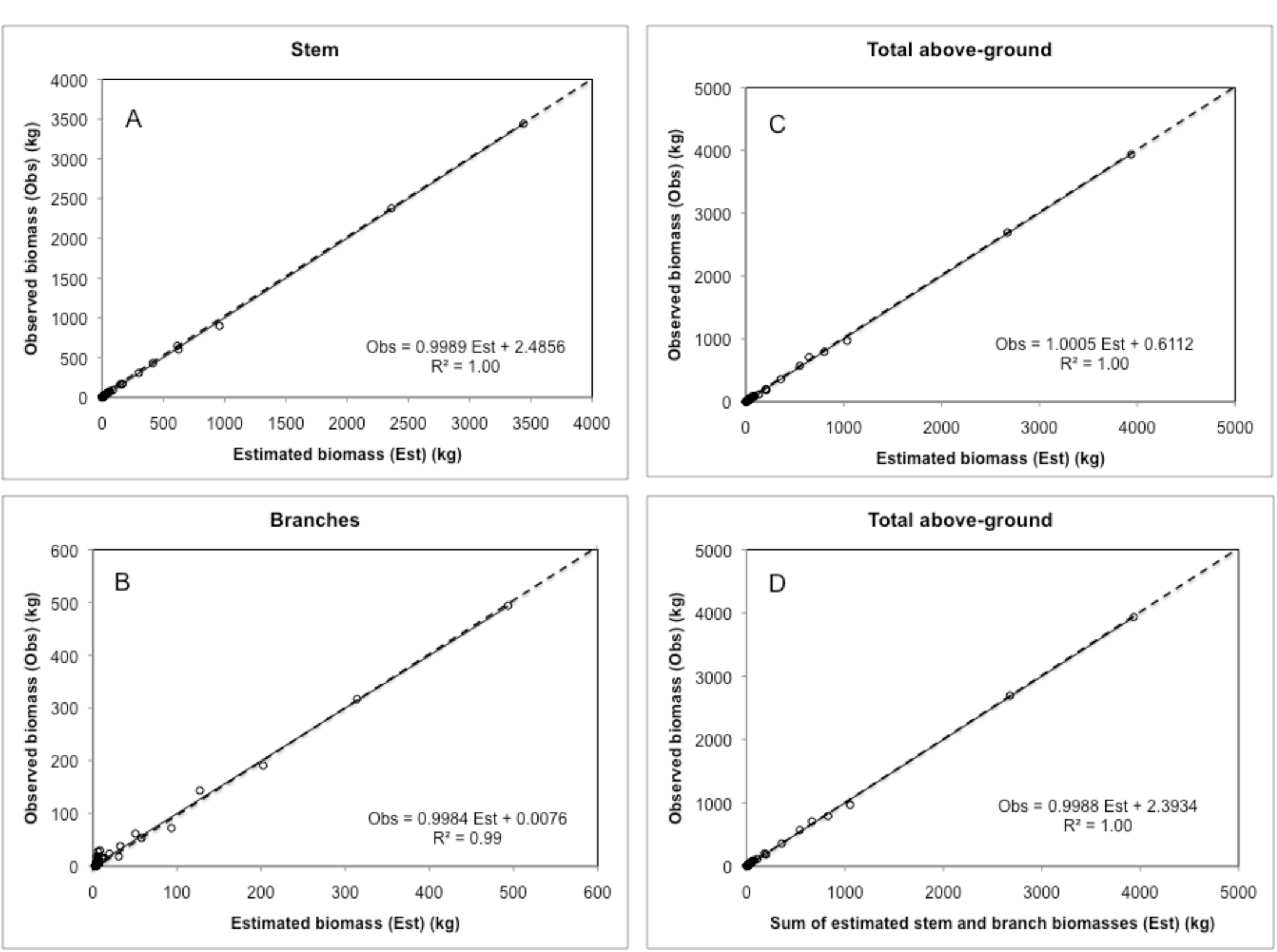
Observed versus estimated biomasses of stem (A), branches (B) and total aboveground, either considered as a whole (C), or as the sum of stem and branches compartments (D): the similarity of the regression lines with the 1:1 dashed lines is obvious.

Stem and branch bark biomasses appeared proportional to stem and branch biomasses, with proportionality coefficients equal to 0.0689 (SE = 0.0004) and 0.1288 (SE = 0.0032) respectively. Stem and branch wood biomasses can then be derived by subtracting stem and branch bark biomasses from total stem and total branch biomasses respectively.

##### belowground

Biomass equations were established for the following belowground tree compartments: coarse roots, small roots, fine roots and whole roots (= coarse + small + fine roots)^11^. Equation (5) could not be fitted to belowground data, maybe due to an excessive number of parameters. The reduced model *W* = (*a* + *b* ^∗^ *Age*) ^∗^ (*D*^2^ ^∗^ *Ht*)^*e*^, with only 3 parameters, was successfully fitted to the data from each compartment, but biased estimations were observed for small and fine roots. The following alternative model gave better results:

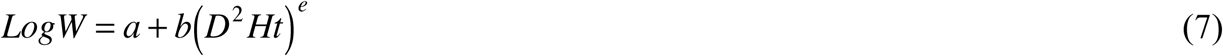

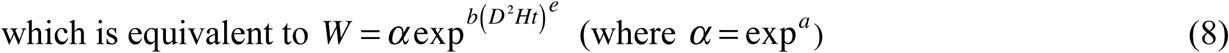

The simultaneous estimation of the parameters of Eq. 7 for each root compartment, with starting values obtained after a separate fitting of each equation, produced the following results (Table 5):

**Table 5.** Estimated values of the parameters of Eq. 7, with their standard errors (in parenthesis), for each root category (coarse, small and fine roots) and for the whole roots compartment (*all parameters significant at the 0.05 level or more*).

| Parameters | Belowground tree compartments |  |  |  |
| --- | --- | --- | --- | --- |
|  | Coarse roots | Small roots | Fine roots | Whole roots |
| *a* | -15.827<br>(5.002) | -8.472<br>(1.673) | -6.661<br>(1.104) | -12.868<br>(3.843) |
| *b* | 12.096<br>(4.808) | 3.743<br>(1.475) | 2.332<br>(0.913) | 9.638<br>(3.650) |
| *e* | 0.05204<br>(0.01541) | 0.10131<br>(0.02411) | 0.12639<br>(0.02690) | 0.05948<br>(0.01614) |

The observation of the residuals of Equation 7 did not reveal any bias in the biomass estimation of each belowground tree compartment, nor any relation with tree variables not already entered in the equation, only a weak dependence with the social status of trees. Random effects linked to social status and attached to the parameter *a*, were then added in Equation 7 fitted in this case with *nlme* (R). The comparison of the estimations obtained with *nls* or *nlme* – with or without consideration of a correlation effect between tree compartments – did not show a significantly better fitting when adding random effects in Equation 7; then, it was decided to keep the simpler model with only fixed effects considered (Table 5).

The estimation of the biomass of the different belowground tree compartments was then obtained by using Equation 8 and multiplying the estimated values obtained by the correction factor
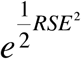
(Flewelling, 1981), where *RSE* is the residual standard error of Equation 7 (*RSE* = 0.3534).

The estimated biomass of the different belowground tree compartments compared favorably with the corresponding observed biomass values (Figures 4A, 4B, 4C, 4D), except for the larger tree of the sample (tree # 35) for which the biomass of coarse roots and of the whole root system seemed overestimated^12^ (Figs 4A, 4D). Moreover, the sum of the estimated biomasses of each root category fitted relatively well the total belowground biomass estimated as a whole (Figure 4D).

**Figure 4.**
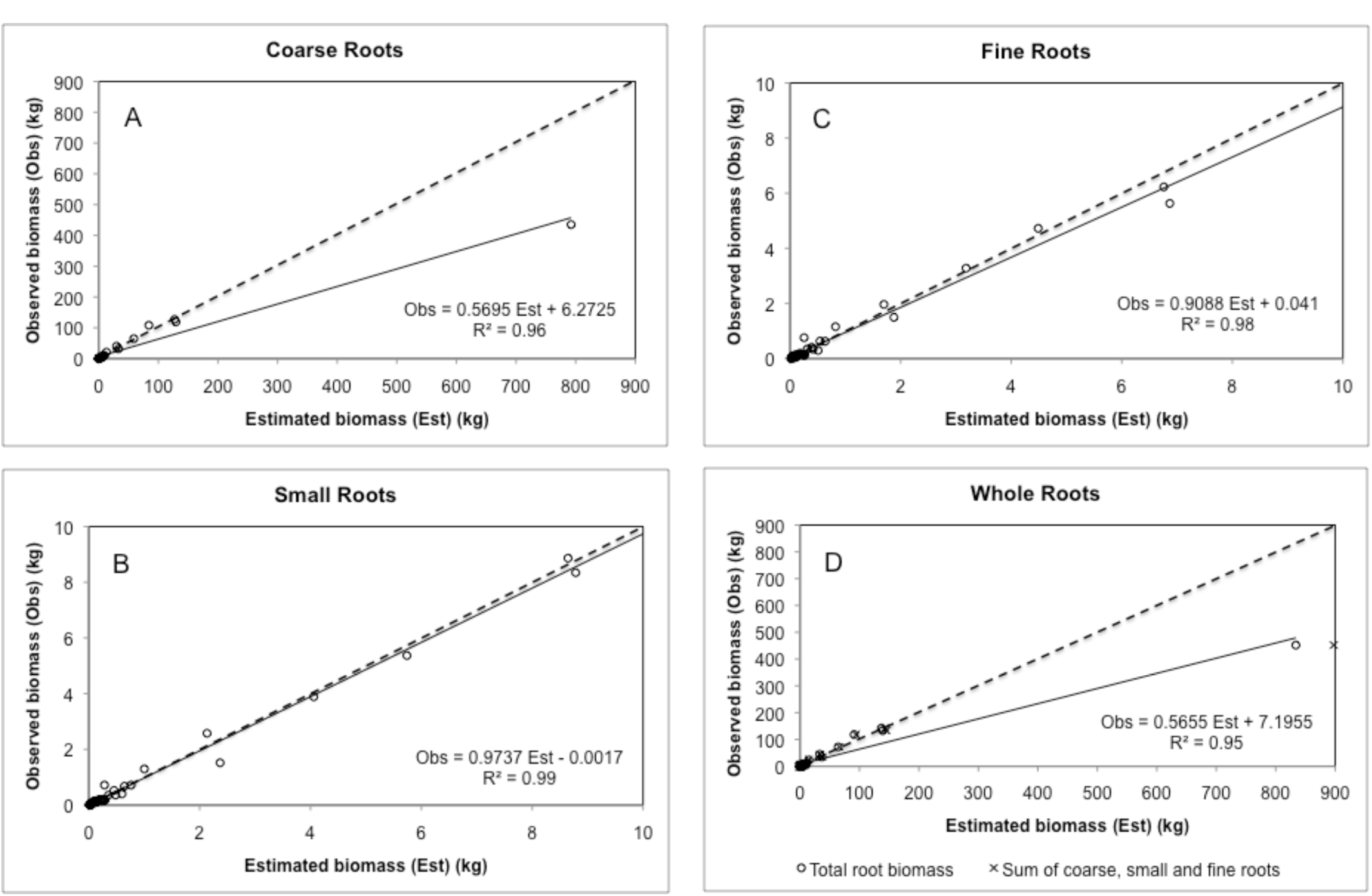
Observed versus estimated biomasses of coarse roots (A), small roots (B), fine roots (C) and of total belowground, either considered as a whole or as the sum of the different root compartments (D) : there is a fairly great similarity of the regression lines with the 1:1 dashed lines for small and fine roots, but not for coarse roots because of an outlier (tree #35, see text).

##### foliage

The model described by Eq. 5 for aerial tree compartments was tested for foliage biomass (*W_F_*). Several parameters in Eq. 5 were not significant and the following reduced model *W* = *a* ^∗^ (*D*^2^*HT*)^*e*+*f*^∗^*Age*^ was then fitted to foliage biomass data after a log-transformation of both sides of the equation. An additional variable (*RCL* = relative crown length) was introduced in the model after observation of the residuals of the preceding equation. The final model (Eq. 9) was fitted using non-linear least-square regression (*nlme*) to take account of the yearly random effects further detected and which were supported by the parameter “a”:

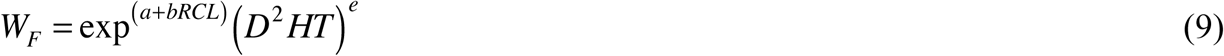

The estimated values of the parameters of Eq. 9, which correspond to fixed effects, and the random effects associated with the parameter “*a*” are listed in Table 6.

**Table 6.** Estimated values and standard errors of the parameters of Eq. 9 linking foliage biomass (*W_F_*, *kg*) to tree dbh (*D*, *cm*), height (*HT*, *m*) and relative crown length (*RCL*), with fixed and random effects distinguished for the parameter “a”.

| Parameter | Fixed effects |  | Random effects (Year) |  |  |  |  |  |  |
| --- | --- | --- | --- | --- | --- | --- | --- | --- | --- |
|  | Estimate | Std error | 1996 | 1997 | 1999 | 2000 | 2001 | 2002 | 2003 |
| a | -7.870 | 0.534 | 0.475 | 0.535 | -0.416 | 0.066 | 0.089 | -0.687 | -0.063 |
| b | 3.781 | 0.397 |  |  |  |  |  |  |  |
| e | 0.813 | 0.057 |  |  |  |  |  |  |  |

The estimated foliage biomasses of sampled trees were in relatively good accordance with the observed values (Fig. 5), and no bias could be detected. Moreover, the examination of the residuals did not reveal any additional effect of the sampling years nor of tree sampling characteristics (forest plot or tree social class belonging).

**Figure 5.**
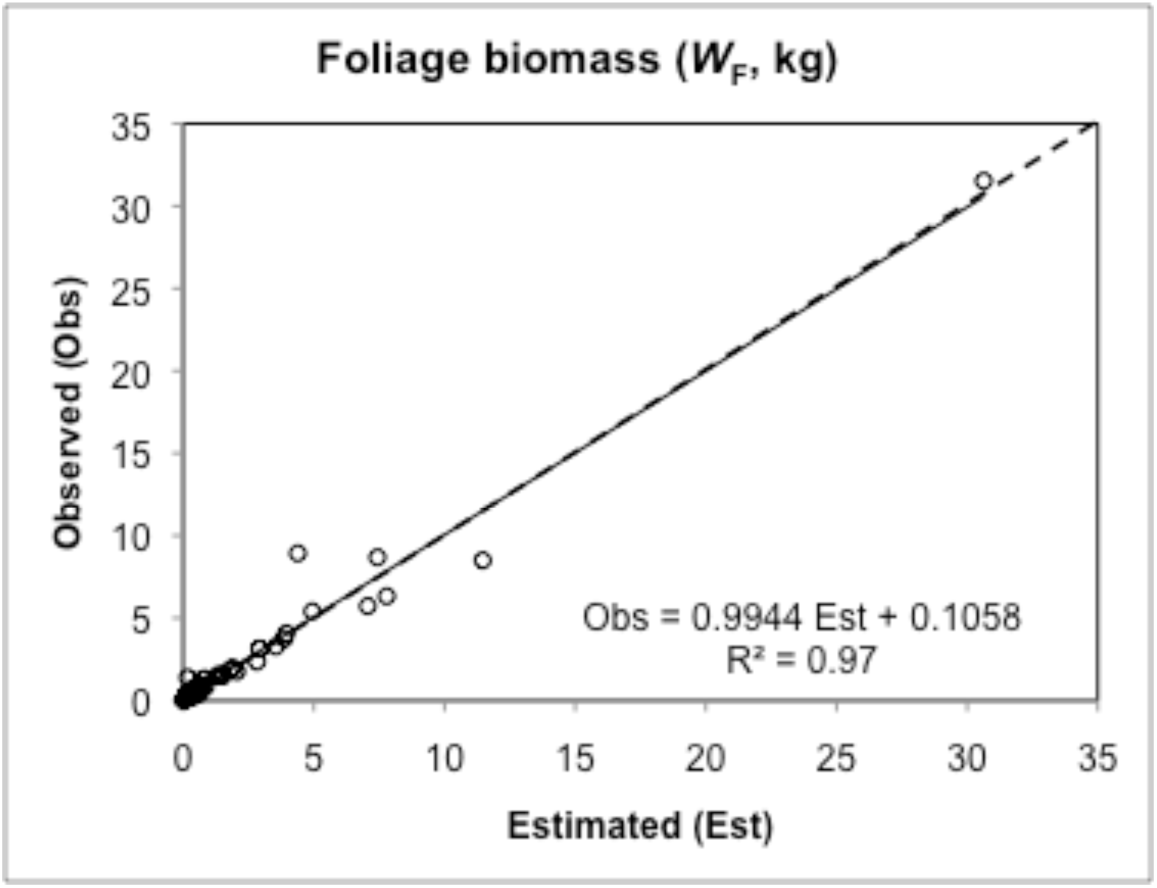
Observed versus estimated (Eq. 9, Table 6) foliage biomasses of 59 sampled trees of various ages in Hesse forest (NE France). (The similarity of the regression line with the 1:1 dashed line is obvious).

Tree leaf area (*LA*) appeared proportional to foliage biomass (*W_F_*), and then a linear relation was fitted using *lme* (R), as the slope coefficient (*k*) in the relation *LA* = *kW_F_* presented random effects linked to the year of tree sampling (Table 7). The estimated leaf area values were in good accordance with the observed ones (Fig. 6), and no other significant effect could be detected.

**Table 7.**
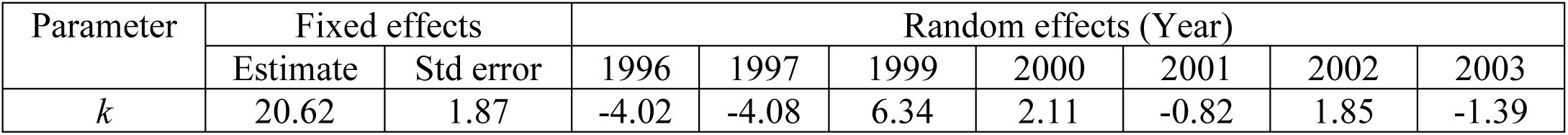
Estimated value of the slope coefficient *k* linking leaf area (*LA*) to foliage biomass (*W_F_*), with fixed and random effects distinguished.

**Figure 6.**
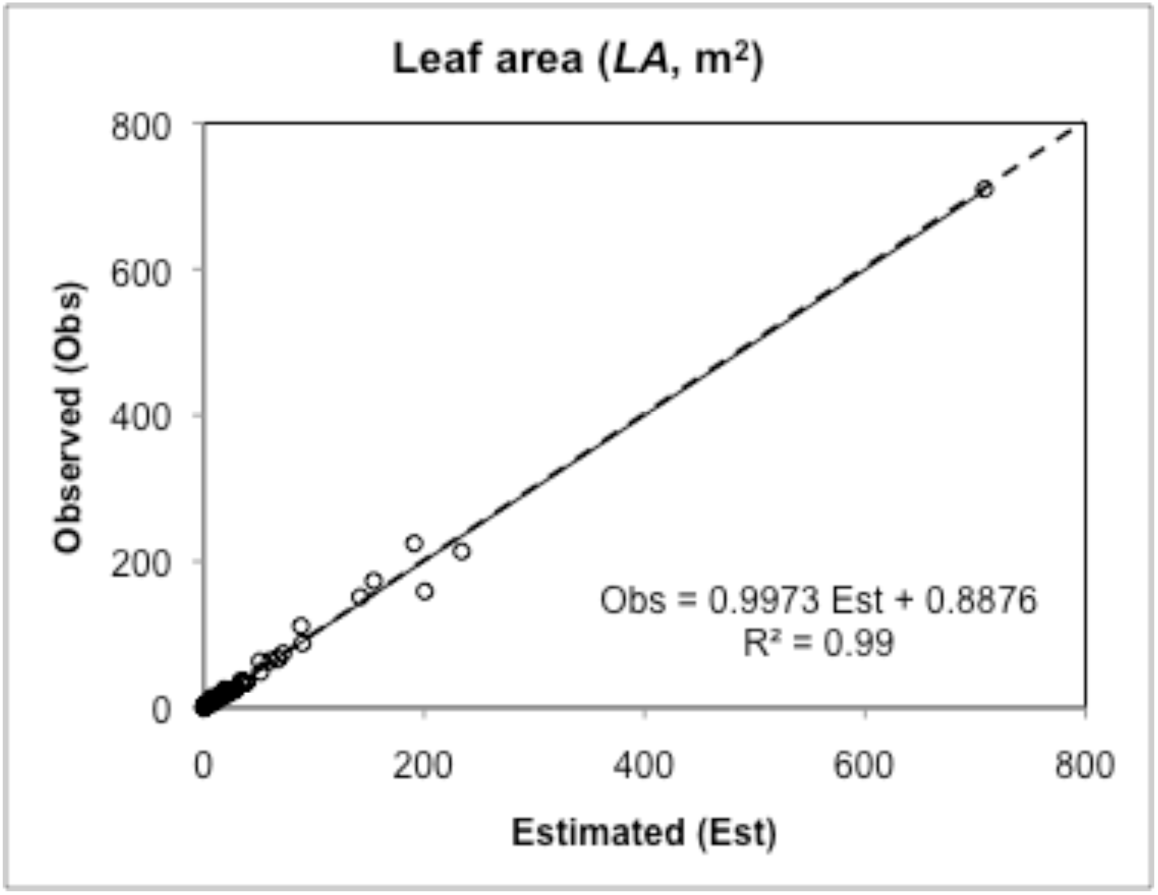
Observed versus estimated leaf areas of 59 sampled trees of various ages in Hesse forest (NE France) obtained from the linear relation between leaf area and foliage biomass (Table 7). (The similarity of the regression line with the 1:1 dashed line is obvious).

The mean specific leaf area (SLA, cm^2^ g^−1^) of the sampled trees was obtained by multiplying the slope coefficient *k* in the preceding relation
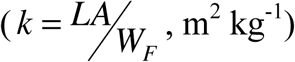
by 10, leading to the following value: SLA = 206.2 cm^2^ g^−1^.

##### biomass distribution

The biomass equations fitted were used to analyze the biomass distribution in above and belowground compartments in relation to tree dimensions (diameter and height) and age. As height (*HT*) is correlated with diameter (*D*) at a given age, a height-diameter equation with age-dependent parameters was fitted to sampled trees data. The following power model was retained after fitting separate equations to data from different age classes and analyzing the variation of equation parameters with age:

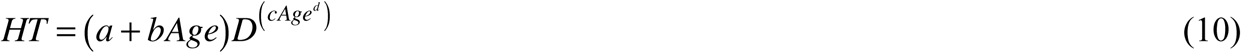

In this equation, the parameter *a* appeared to be dependent on forest plot. Then, Eq. 10 was fitted with *nlme* (R), fixed and random effects appearing in Table 8. Plotting residuals against fitted values did not reveal any bias.

**Table 8.** Estimated values of the parameters of Eq. 10 linking tree height (*HT*) to diameter at breast height (*D*) and age (*Age*), with fixed and random effects distinguished.

| Parameter | Fixed effects |  | Random effects (forest plot) |  |  |  |  |
| --- | --- | --- | --- | --- | --- | --- | --- |
|  | Estimate | Std error | P214 | P215 | P217 | P220 | P222 |
| a | 5.3402 | 0.6915 | 0.5316 | -1.5525 | 0.5495 | -0.5049 | 0.9762 |
| b | 0.0041 | 0.0157 |  |  |  |  |  |
| c | 0.1748 | 0.0776 |  |  |  |  |  |
| d | 0.1987 | 0.1214 |  |  |  |  |  |

Biomass distribution in trees was then represented in relation to diameter at breast height and age, replacing tree height (*HT*) in biomass equations by its estimation obtained from Eq. 10. In Fig. 7, the contributions of the main above and belowground compartments to tree biomass is represented, while in Fig. 8 the contributions of the different root compartments (coarse, small and fine roots) to total belowground biomass are represented.

**Figure 7.**
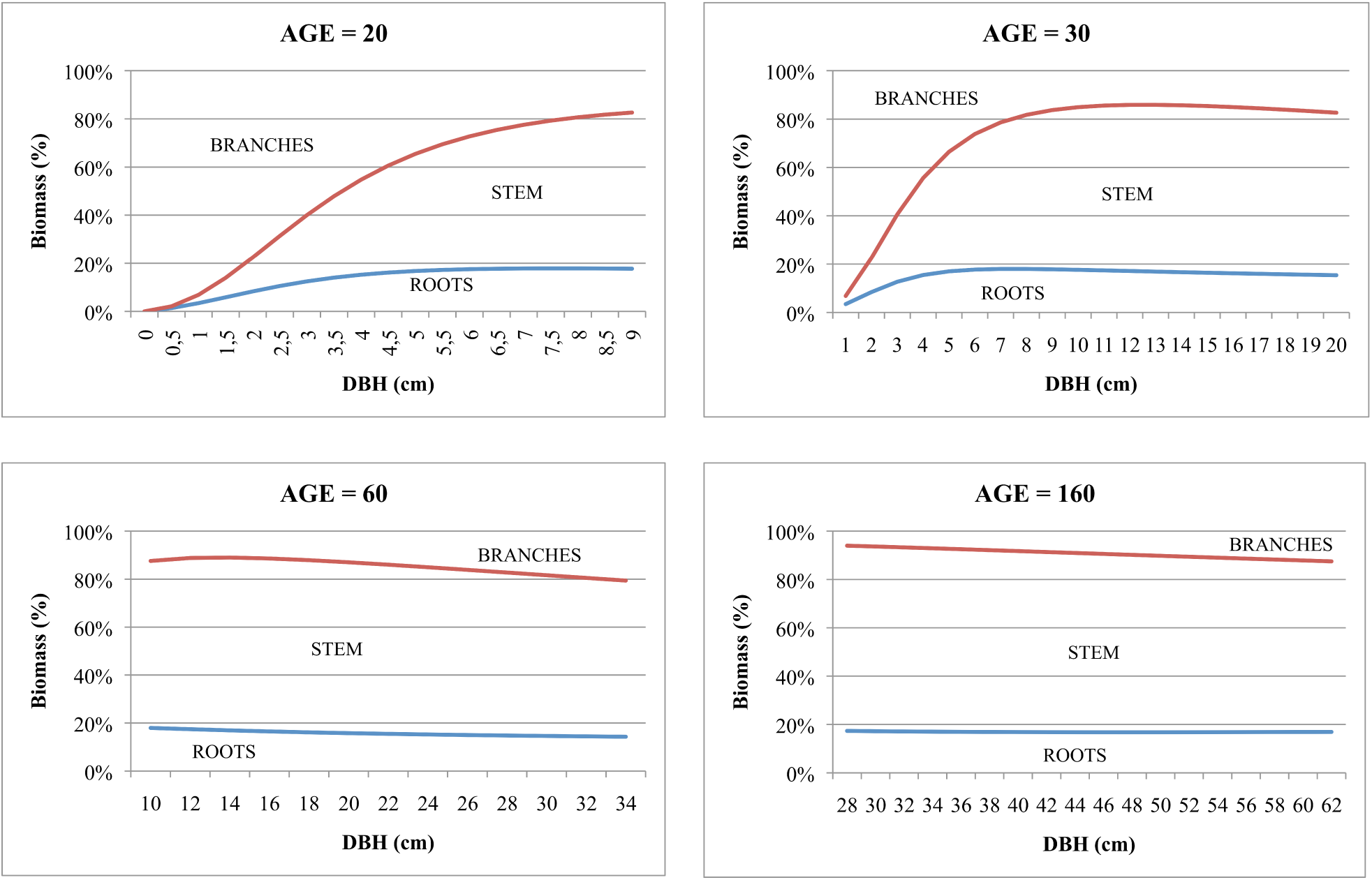
Distribution of the biomasses of the main tree compartments – stem, branches and roots – in relation to tree diameter D (cm) and age.

**Figure 8.**
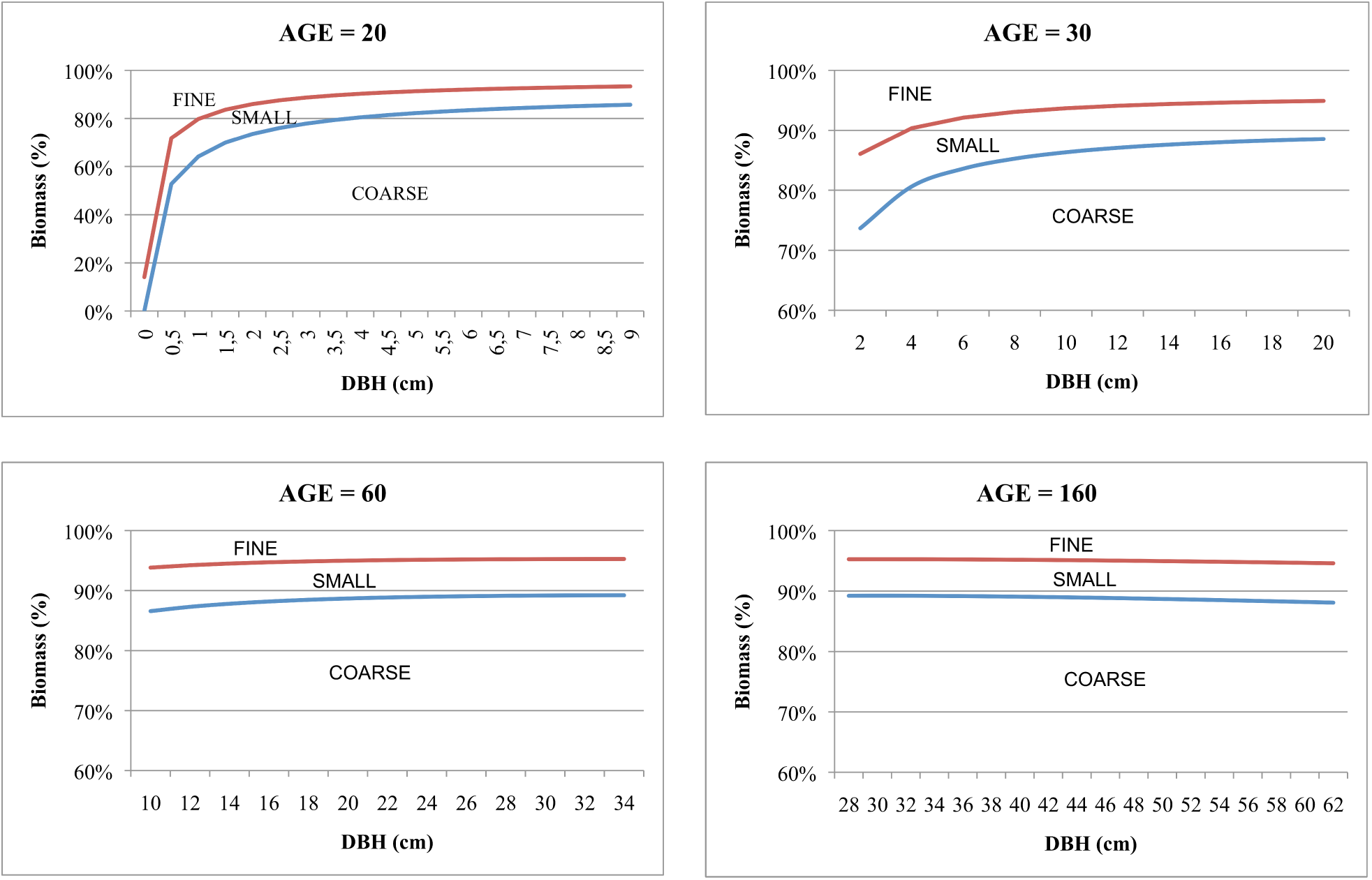
Distribution of the biomasses of the different root compartments – coarse, small and fine roots – in relation to tree dbh (cm) and age.

Tree stem appears as the main tree compartment, representing more than 60% of the tree biomass, except maybe in the young ages where branch biomass seems to exceed that of the stem for the smallest trees. Roots contribute to tree biomass at a relatively constant rate across ages, the mean root-shoot ratio^13^ being equal to 0.23 for the whole sub-sample of trees, with slightly higher values (0.32) for the youngest trees of the sample aged about 20.

Coarse roots contribute to about 90% of total belowground biomass for the largest trees, independently of age class. This contribution tends to decrease with decreasing size of trees when trees are young, for the benefit of small and fine roots compartments.

### 3.3 Biomass increment

#### 3.3.1 Biomass increment of aerial compartments

The following model, derived from the model of Eq. 6, was retained to represent the variations of the biomass increment of aboveground tree compartments (Δ*W*) with tree leaf area (*LA*), age (*Age*) and foliar density (*DSF*)^14^ :

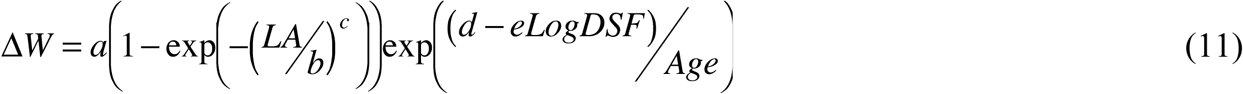

Equation 11 was simultaneously fitted to each aboveground tree compartment (stem and branches) and to total aboveground. As the biomass increment of the aboveground tree compartments (stem and branches) appeared proportional to total aboveground biomass increment, only the parameter *a* in the above equation varied with tree compartments, and Eq 11 was re-written and fitted as follows:

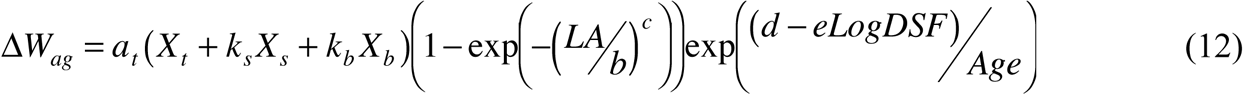

In Eq.12, X_t_, X_s_ and X_b_ are dummy variables taking the value “1” for total aboveground, stem or branch compartments respectively (“0”, otherwise); *a_t_* is the multiplicative parameter for total aboveground biomass increment, *k_s_* and *k_b_* are the multiplicative factors of *a_t_* for stem and branch biomass increments respectively; *b*, *c*, *d* and *e* are common parameters.

To estimate the parameters of Eq. 12, in which Δ*W_ag_* stands for the biomass increment of either aboveground tree compartment (stem, branches or total aboveground), a weighted non-linear mixed-effects regression was performed (*nlme*, R) with random effects due to forest plot belonging supported by the parameter *a_t_.* The introduction of correlations among the parameters, in relation with the inter-dependence of tree compartments, was not retained as it did not change much the estimated values of the parameters.

The estimated values of the parameters of Eq. 12, with their standard errors, are listed in Table 9. All parameters were significant at the 0.001 level. Random effects associated with forest plot belonging (P214 to P222) are also listed.

**Table 9.** Estimated values of the parameters of Eq. 12, with fixed and random effects distinguished for the multiplicative parameter which takes the values *a_t_*, *a_t_k_s_* and *a_t_k_b_* for total aboveground, stem and branch biomass increments respectively.

| Parameters<br>(Eq. 11) | Fixed effects |  | Random effects (forest plot) |  |  |  |  |
| --- | --- | --- | --- | --- | --- | --- | --- |
|  | Estimate | Std error | P214 | P215 | P217 | P220 | P222 |
| $a_t$ | 68.322 | 13.464 | -34.928 | 42.618 | 3.471 | -20.936 | 9.775 |
| $k_s$ | 0.7587 | 0.0467 | 0.0416 | -0.1594 | 0.0699 | 0.0982 | -0.0504 |
| $k_b$ | 0.2413 | 0.0441 | -0.0391 | 0.1580 | -0.0713 | -0.0978 | 0.0503 |
| $b$ | 344.71 | 15.238 | | | | | |
| $c$ | 1.5826 | 0.0380 | | | | | |
| $d$ | 88.577 | 7.347 | | | | | |
| $e$ | 28.290 | 3.442 | | | | | |

The observation of the residuals of Eq. 12 did not reveal any bias in the biomass increment estimations, and no other significant effect on the biomass increment of the different tree compartments could be detected, especially in relation with the social status of trees or the year of tree sampling. The observed and estimated values of the biomass increments of the aboveground tree compartments of the sampled trees in Hesse forest compared relatively well (Figs. 9A, 9B, 9C). Moreover, the sum of the estimated biomass increments of each aboveground tree compartment fitted relatively well the aboveground biomass increment estimated as a whole (Figure 9D).

**Figure 9.**
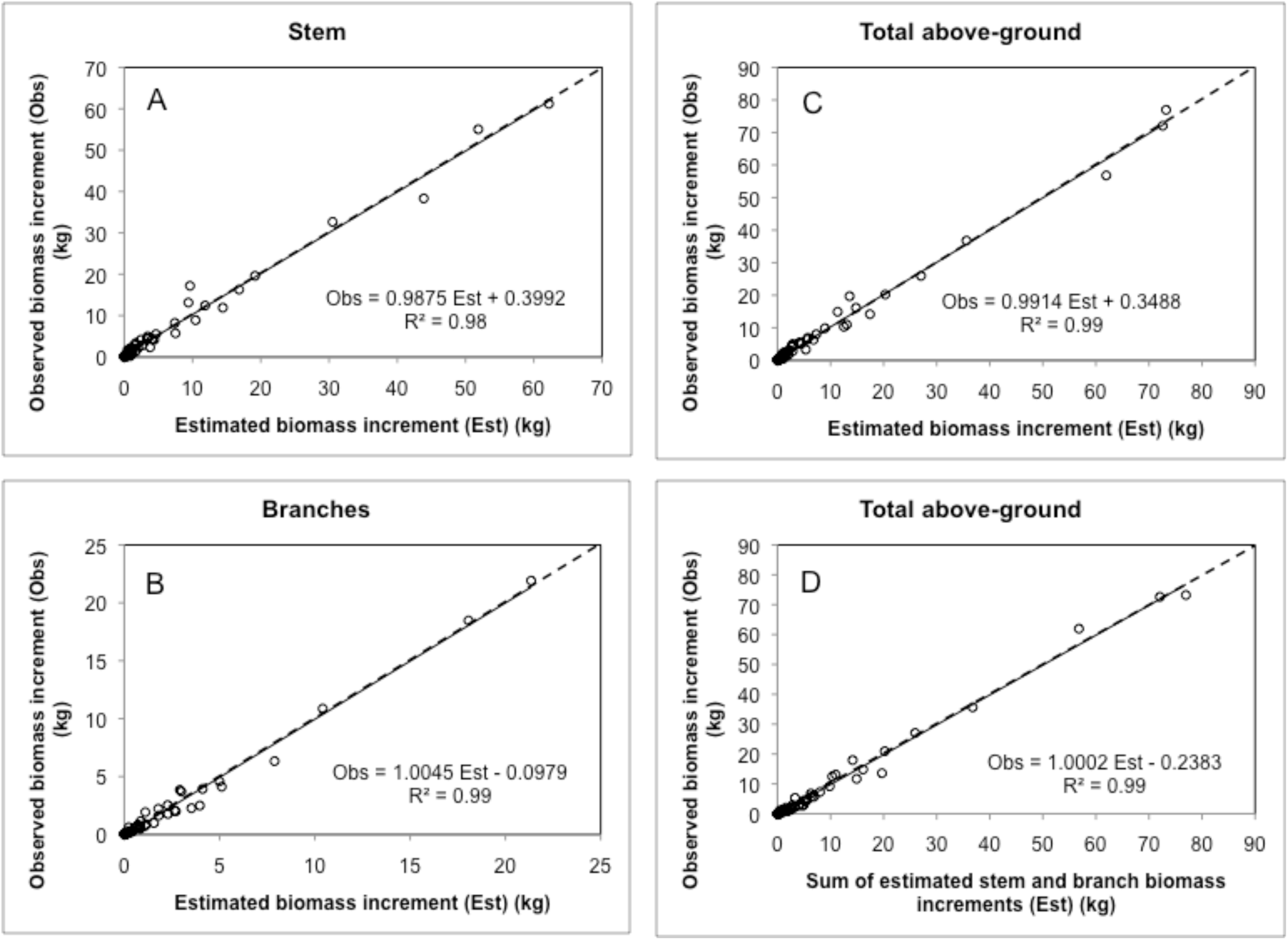
Observed versus estimated annual biomass increments of stem (A), branches (B) and total aboveground, either considered as a whole (C) or as the sum of stem and branch compartments (D): the similarity of the regression lines with the 1:1 dashed lines is obvious in each case.

The equations, specific of each aboveground tree compartment, could be derived from Eq. 12:

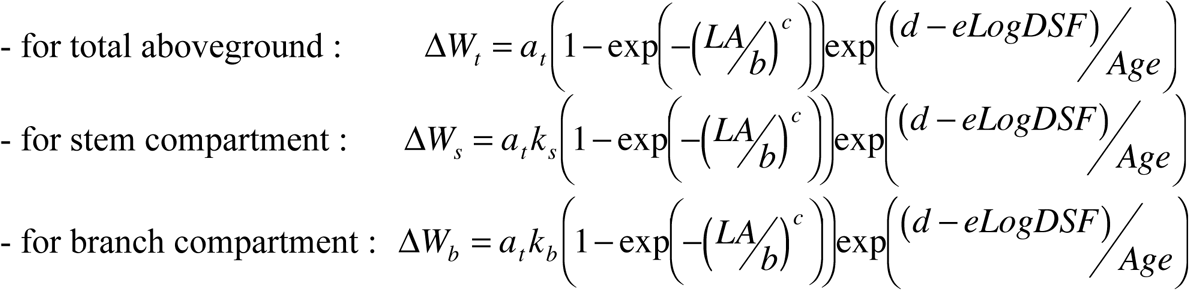

#### 3.3.2 Root biomass increment

The model described by Eq.11 allowed the representation of the variations of the biomass increments of the belowground tree compartments (coarse roots, small roots and total root system), which appeared also proportional. Then, Eq. 12 was adapted for belowground tree compartments and fitted in the following form:

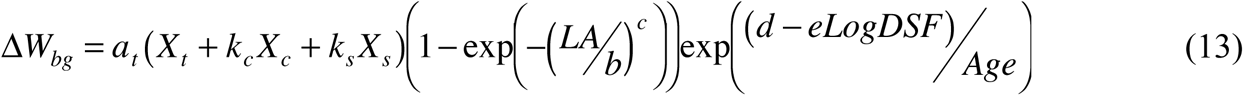

In Eq. 13, X_t_, X_c_ and X_s_ are dummy variables taking the value “1” for total belowground, coarse roots and small roots compartments respectively (“0”, otherwise); *a_t_* is the multiplicative parameter for total belowground biomass increment, *k_c_* and *k_s_* are the multiplicative factors of *a_t_* for coarse and small roots respectively; *b*, *c*, *d* and *e* are common parameters for the different root compartments.

A non-linear regression model was fitted to Eq. 13 using *nls* (R). Weights and correlations were introduced in the model as before, but seemed to generate biased estimations and then were not retained in the fitting process. Furthermore, no significant random effect – in relation with tree location (forest plot) or tree social class belonging – could be detected. Additionally, no significant effect of the year of tree sampling could be detected.

The estimated values of the parameters of Eq. 13, with their standard errors, are listed in Table 10. All parameters were significant at the 0.001 level.

**Table 10.** Estimated values of the parameters of Eq. 13 with their standard errors; the multiplicative parameter *a* of Eq. 11 takes here the values *a_t_*, *a_t_k_c_* and *a_t_k_s_* for total belowground, coarse roots and small roots biomass increments respectively.

| Characteristic | Parameters of Eq. 13 |  |  |  |  |  |  |
| --- | --- | --- | --- | --- | --- | --- | --- |
| | $a_t$ | $k_c$ | $k_s$ | $b$ | $c$ | $d$ | $e$ |
| Estimate | 7.2673 | 0.9507 | 0.0493 | 236.958 | 1.9492 | 117.565 | 34.570 |
| Stand. Error | 0.534 | 0.0170 | 0.0124 | 7.120 | 0.0425 | 14.201 | 7.974 |

The observed and estimated values of the biomass increments of the belowground tree compartments of the sampled trees compared relatively well (Fig. 10A, 10B, 10C). Moreover, the sum of the estimated biomass increments of each belowground tree compartment equaled the belowground biomass increment estimated as a whole (Fig. 10D).

**Figure 10.**
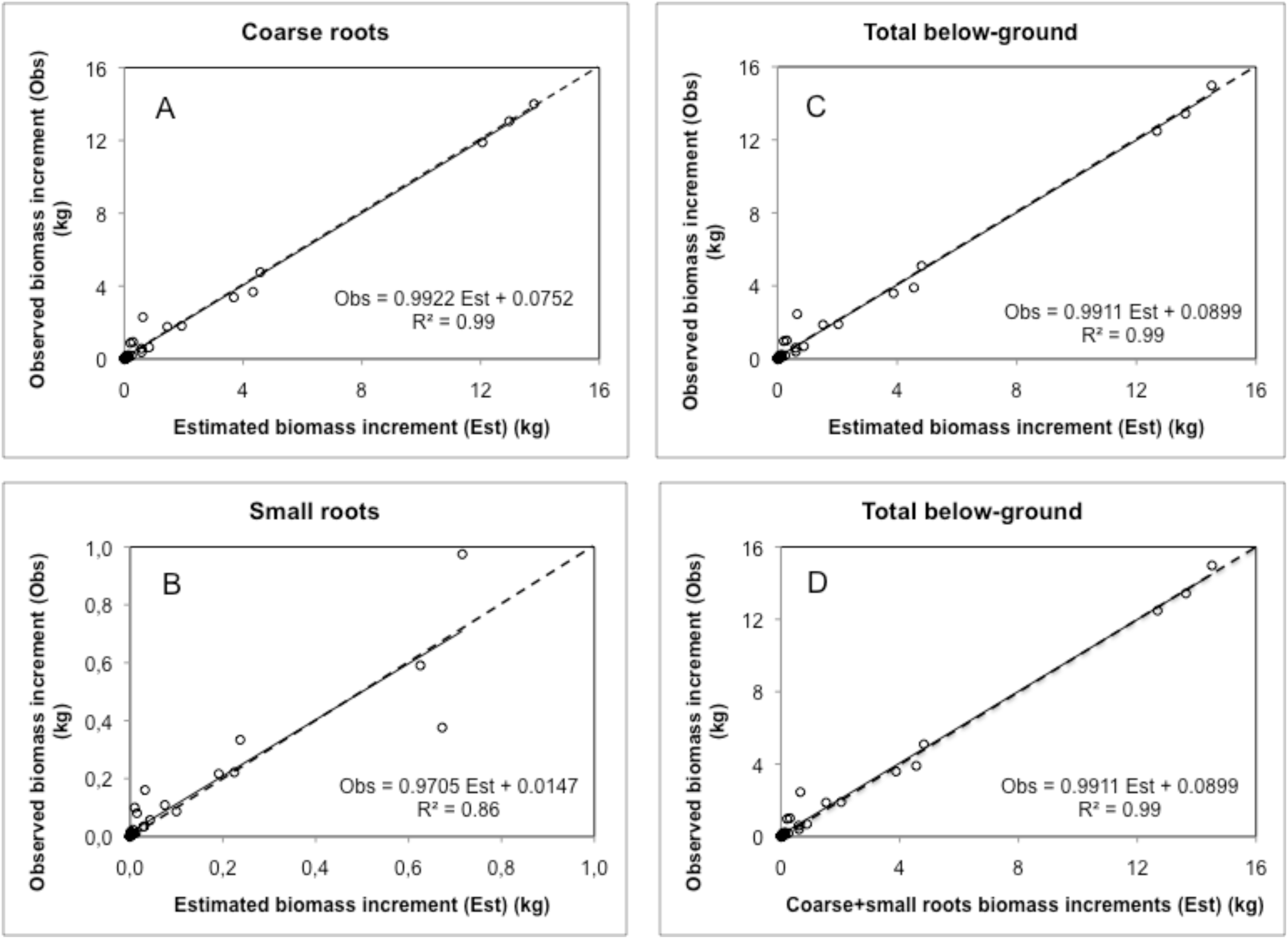
Observed versus estimated annual biomass increments of coarse roots (A), small roots (B) and total belowground, either considered as a whole (C) or as the sum of coarse and small roots (D): the similarity of the regression lines with the 1:1 dashed lines is obvious in each case.

The equations specific of each belowground tree compartment could be derived from Eq. 13 with the parameters values listed in Table 10:

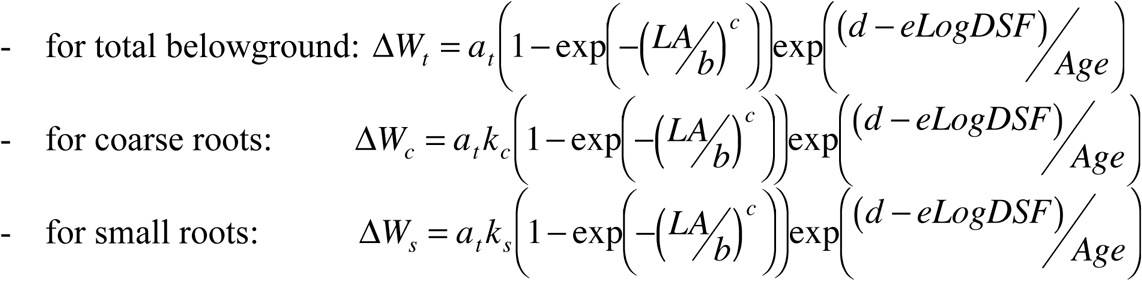

### 3.4 Biomass allocation

In order to analyze the variations of the distribution of biomass increment in trees with age, leaf area (LA) and density of foliage (*DSF*) – from which depends tree growth – *LA* and *DSF* were related to tree age (*Age*). The following relations could be established, using non-linear and linear regressions for *LA* and *DSF*, respectively:

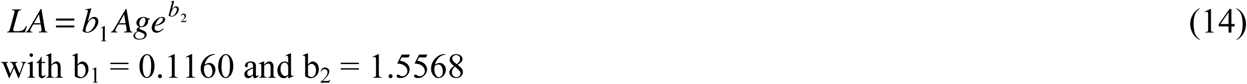

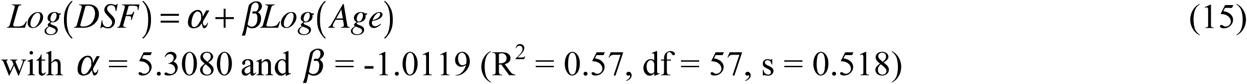

The distribution of the biomass increment in trees was then represented in relation to tree age, replacing leaf area (*LA*) and density of foliage (*DSF*) in biomass increment equations by their estimations obtained from Eq. 14 and Eq. 15 respectively^15^. In Fig. 11, the contributions of the main above and belowground compartments to tree biomass increment are represented: biomass increment appears preferentially allocated to the stem (more than 60%), then to branches (about 20%) and roots (less than 20%). With regard to the root compartment, coarse roots appear to contribute to 95% of total belowground biomass increment while small roots contribute to 5% only, independently of tree age^16^.

**Figure 11.**
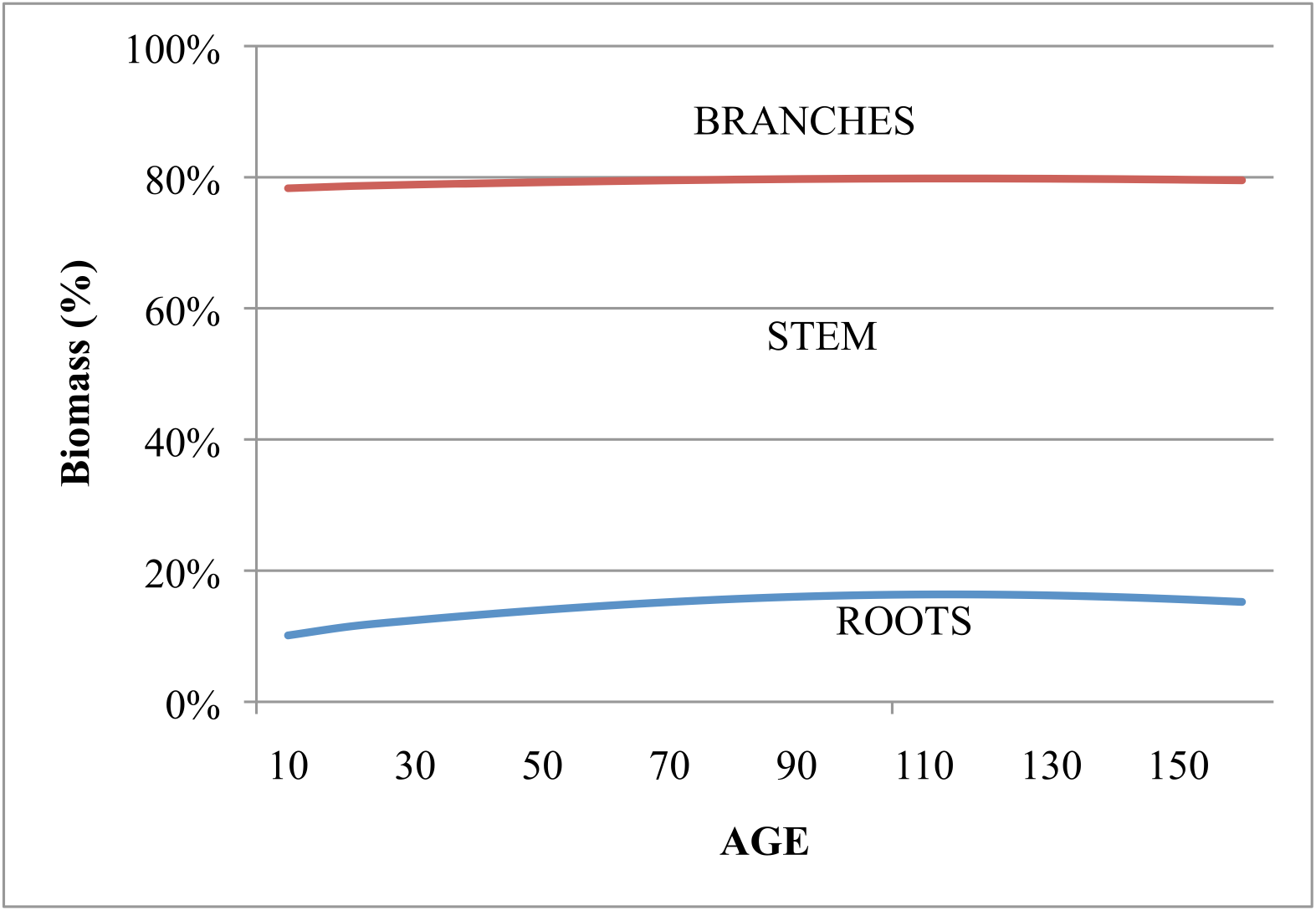
Distribution of the biomass increments among the main tree compartments, in relation to tree age.

#### 3.5 Stem volume increment and growth efficiency

The model developed to represent the variations of the annual biomass increment of aboveground tree compartments (Eq. 11) was retained to represent the variations of bole (or stem) annual volume increment (*BI*), that is:

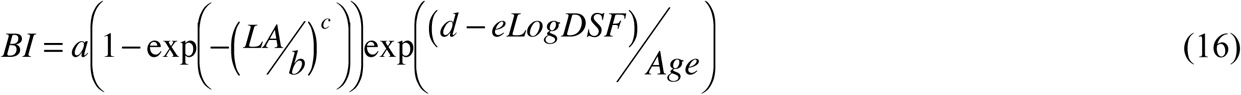

To estimate the parameters of Eq. 16, a non-linear mixed-effects regression was performed (*nlme*, R) with random effects due to forest plot belonging supported by the parameters *b* and d. The estimated values of the parameters of Eq. 16, with their standard errors, are listed in Table 11. Random effects associated with forest plot belonging (P214 to P222) are also listed for *b* and *d* parameters.

**Table 11.** Estimated values of the parameters of Eq. 16 linking bole volume increment *BI* to leaf area (*LA*), density of foliage (*DSF*) and age (*Age*), with fixed and random effects distinguished for the parameters *b* and *d.*

| Parameters<br>(Eq. 16) | Fixed effects |  | Random effects (forest plot) |  |  |  |  |
| --- | --- | --- | --- | --- | --- | --- | --- |
|  | Estimate | Std error | P214 | P215 | P217 | P220 | P222 |
| *a* | 65.8972 | 5.5281 |  |  |  |  |  |
| *b* | 361.8440 | 65.2292 | 170.811 | 32.418 | -55.867 | -18.408 | -128.954 |
| *c* | 1.5018 | 0.05293 |  |  |  |  |  |
| *d* | 114.4202 | 18.9182 | 28.235 | 5.359 | -9.235 | -3.043 | -21.316 |
| *e* | 35.3697 | 7.6299 |  |  |  |  |  |

No other significant effect on *BI* could be detected, including the social status of trees and the year of tree sampling, and the fitted values of *BI* appeared in good agreement with the observed ones (Fig. 12).

**Figure 12.**
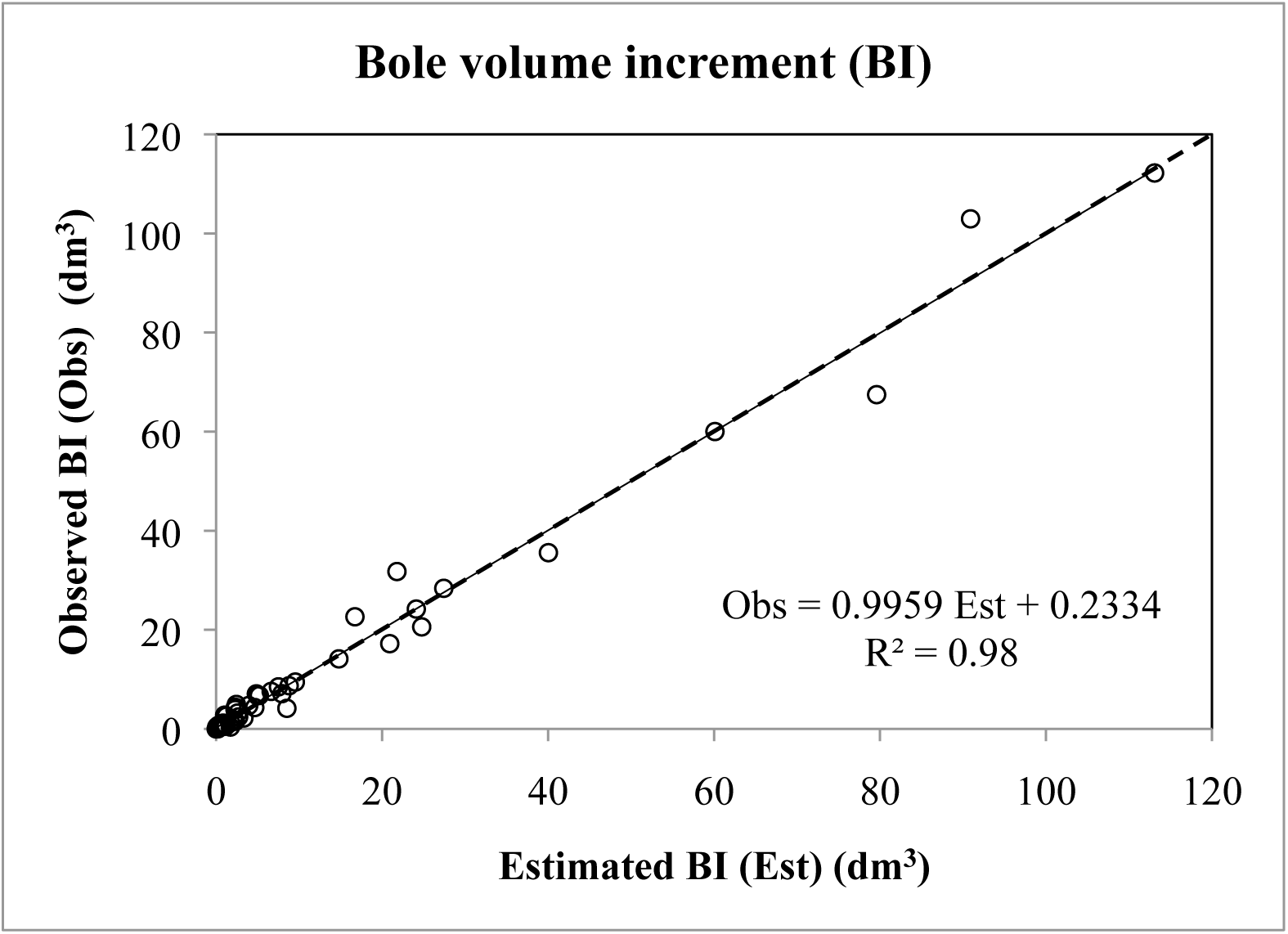
Comparison of the observed and estimated (Eq. 16) annual bole volume increments (BI) of trees for the tree sample of Hesse forest (NE France). (The similarity of the regression line with the 1:1 dashed line is obvious)

Growth efficiency (*GE*) of trees, defined as stem annual volume increment per unit of leaf area, was estimated by dividing the *BI* model (Eq. 16) predictions by their corresponding observed leaf areas, as was done by Hofmayer et al (2010) and DeRose and Seymour (2009). The estimated values of *GE* fitted relatively well the observed ones (Fig. 13A).

**Figure 13.**
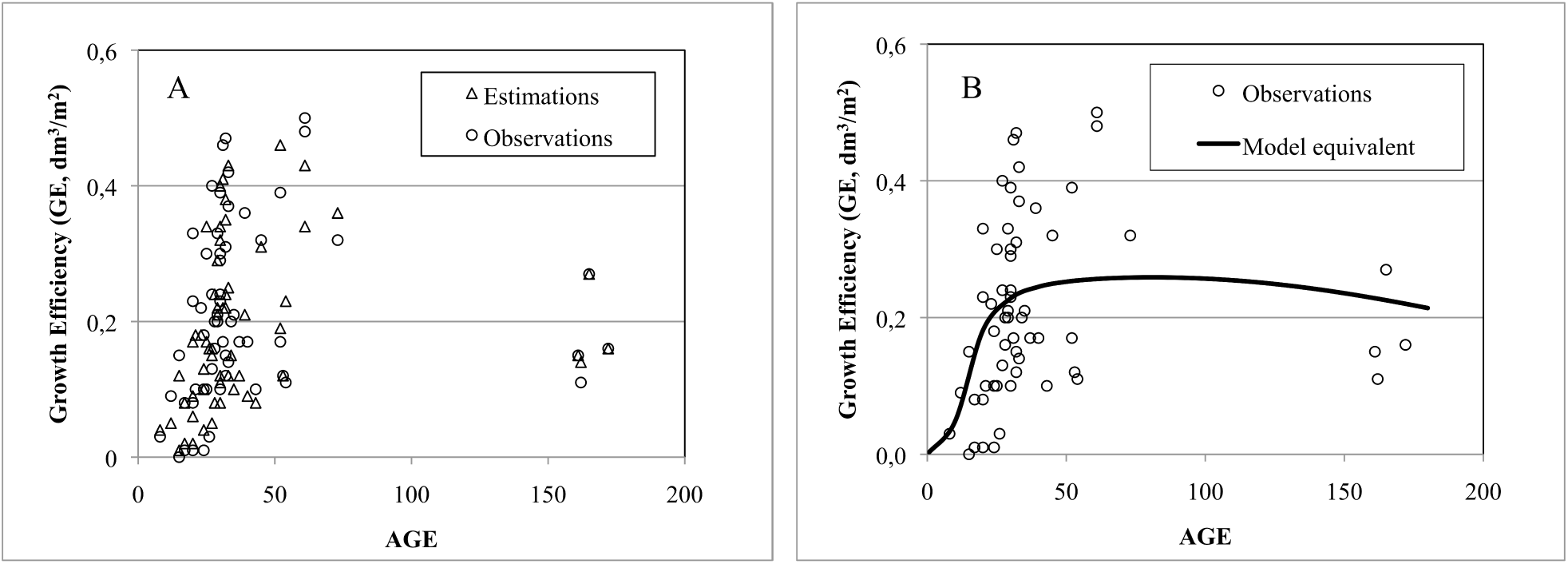
Observed stem volume growth efficiency (*GE*) for the tree sample of Hesse forest versus estimated values obtained from Eq. 16, using either observed (A) or estimated (B) LA and DSF values (*LA* and *DSF* estimates obtained from Eq.14 and Eq.15 respectively).

The simulated *GE* values, calculated by replacing *LA* and *DSF* values by their estimations obtained from Eq. 13 and Eq.14 respectively, were in good accordance with the trend revealed by plotting observed GE values in relation to age (Fig. 13B). The decrease in GE observed for very young and very old trees appears closely reproduced by the model.

## 4. Discussion

The sampling of more than 60 trees of different social status, and representing the range of ages for a total beech stand rotation, allowed to establish generalized biomass equations for the main aboveground tree compartments (stem, branches and leaves) and for total aboveground biomass. Moreover, a subsample of 40 trees, chosen so as to represent the same range of social status and ages as the main tree sample, allowed to obtain biomass equations for the belowground parts of trees defined in relation to root diameter (coarse, small and fine roots), and for total belowground biomass.

The excavation of the root systems of the sampled beech trees caused the loss of some parts of the root systems, and then equations were established so as to estimate the missing parts of each root system by root category, as was done in a preceding study (Le Goff & Ottorini, 2001); in addition, for establishing these equations in the present study, the distinction was made between horizontal and vertical roots for the largest trees. It seems, however, that for the largest tree of the sample (tree # 35), the belowground biomass was under-estimated, due probably to an incomplete inventory of broken roots during excavation, which resulted in an underestimation of the biomass of missing root parts for that tree.

Due to the large dimensions of the oldest trees in the sample, the biomass of the aboveground tree compartments could not be measured directly for large trees, but via volume and wood density evaluations. In addition, all branches were not measured for these trees, and a stratified sample of branches was used and biomass equations were established to estimate the biomass of non-sampled branches.

### 4.1 Fitting biomass and biomass increment models

Linear and non-linear models were examined to fit biomass and biomass increment data. When linear or non-linear models could equally be used, notably after log-transformation, the non-linear ones revealed better suited when looking at the residuals.

When fitting the relations, a particular attention was brought to the unexplained variation and to the quality of the fitting in order to obtain an eventually better fit. Variables not introduced in the models but characterizing the status of sampled trees – tree social status, year of tree sampling, forest plot belonging – were retained for the analysis of residuals and eventually added in the models as random effects. In this last case, mixed models were fitted using the package *nlme* of *R*. It appeared that such random effects were significant for yearly varying biomass variables (foliar biomass, biomass increment of aboveground tree compartments): in this case, it could be interesting to try to identify these effects, probably related to the yearling varying climate and particularly its consequences on the water availability for trees as it is a limiting factor of the growth of beech (Le Goff & Ottorini 1999; Granier & al, 2008).

Additive models should have been fitted for above and belowground data to ensure that the total tree biomass and biomass increment equals respectively the sums of the biomasses and biomass increments of the different compartments in which they were divided. To simplify the fitting process, multivariate models were used which allowed to obtaining a relatively good agreement for the biomass and biomass increments of the above and belowground tree parts estimated as a whole or as the sum of their constituting compartments. This is in line with the results obtained by Repola (2008, 2009).

### 4.2 Biomass models

The generic model −*W* = *α* + *β* ^∗^ (*D*^2^ ∗ *Ht*)^*γ*^ – already used by Genet & al. (2011a), appeared well suited here to represent the biomass variations of aboveground tree compartments with tree dbh (D) and height (Ht). The constant α was significant only for the branch compartment, as it was also the case for Genet & al. (2011a,b). Moreover, as for Genet & al. (2011b), the parameters *β* and *γ* of the biomass model appeared dependent on tree age (Age) and could be expressed with the same functions whose parameters vary with tree compartment (Eq. 5 and Table 4). However, in our case, the parameter *β* was decreasing with age, independently of the tree compartment considered, while it increased for stem compartment in Genet & al. (2011a,b); furthermore, the parameter *γ* slightly increased with age, from about 1 to 1.1 or 1.15 (stem and overall aerial part respectively) or remained constant, close to 1.5 (branches) in our case, while it appeared independent of age regardless of the compartment considered for Genet & al. (2011a,b).

The parameter *β* decreased linearly with increasing tree age for the stem compartment while it decreased exponentially for the branch compartment. As *γ* is constant for branches, this means that branch biomass is comparatively lower for trees with same diameter and height but older, probably in relation with a smaller crown due to higher crowding conditions. This is not true for the stem as the increase of *γ* with age more than compensates for the decrease of *β* with increasing age: the stem biomass is comparatively higher for trees with same diameter and height but which are older, as the stem of more crowded trees is more cylindrical and has then a higher volume (and biomass) for a given diameter and height.

An exponential model – *W* = *α*exp^*b*(*D*^2^*Ht*)^*e*^^ – appeared better suited for belowground biomass data. In this case, the parameters of the model did not seem to depend on tree age, as it was the case in the model fitted for aboveground biomass data. This result confirms preceding findings (Le Goff & Ottorini, 2001).

While tree diameter and height explain a large part of tree biomass variations, tree age appeared also as an important variable to consider, at least for aboveground tree compartments, as already shown by Genet & al. (2011a,b) or Shaiek & al. (2011). In fact, it can take into account the competitive conditions supported by trees which influence the dimensions of tree crowns and then the branch biomass itself, but also the distribution of the increments on the stem and then stem form (stem tapering) and biomass (Repola, 2009). Tree age in biomass equations may also reflect a possible effect of wood density, which tends to increase with age for beech (Bouriaud et al, 2004). However, no age effect could be detected in belowground biomass equations, agreeing with previously published results (Bond-Lamberty & al, 2002; Le Goff & Ottorini, 2001; Genet et al, 2011a,b): this may be due to a weaker connection between the root system dimensions and tree growing conditions, as tree crowding.

Tree foliage biomass could be expressed with the same model as aboveground tree compartments, and then appeared dependent on tree dbh and height. However, foliage biomass was also dependent on tree crown dimensions, increasing with the relative crown length of the tree (*RCL*), as already observed for birch in Finland (Repola, 2008). For a given dbh and height, foliage biomass increases with crown dimensions with which branch biomass increases, the tree being then able to support more foliage. A year-to-year variation of foliage biomass could also be detected. It has been widely documented that the quantity of foliage may vary significatively from year to year for a given tree or stand, in particular in relation with climatic conditions (Breda & Granier, 1996; Breda, 1999; Le Dantec & al, 2000). In the case of Hesse forest – from which the tree sample of this study came from – year-to-year variations of the stand LAI could be observed, apart from those due to thinning operations (Granier & al, 2008). The close proportional relation observed between tree leaf area and biomass of sampled beech trees allowed to estimate a mean specific leaf area (SLA) value of 206.2 cm^2^ g^−1^ for these trees: Bartelink (1997) observed also such a relationship for beech leaf area, leading to a mean SLA value of 172 cm^2^ g^−1^ for a smaller sample of trees ranging in age from 8 to 59 years, a SLA value comparable to that found in our study.

### 4.3 Biomass distribution in trees

Tree stem represents the main part of tree biomass (between 60 and 80%), except in young ages where the branches may contribute more than the stem to tree biomass in the smallest trees. When trees are ageing, the branch contribution to tree biomass tends to decrease to less than 20%, and relatively more for smaller trees (Fig. 7). Regarding diameter as an indicator of tree status in stands at a given age, this means that dominance, or tree competitive status, affects the amount of branch biomass (Bartelink, 1997): with increasing inter-tree competition, a lower fraction of tree biomass is represented in branches, except maybe for suppressed trees in young stands (Fig. 1). Stem and branch biomass data in our study appear consistent with those obtained by Bartelink (1997), when related to tree diameter (Fig. 14).

**Figure 14.**
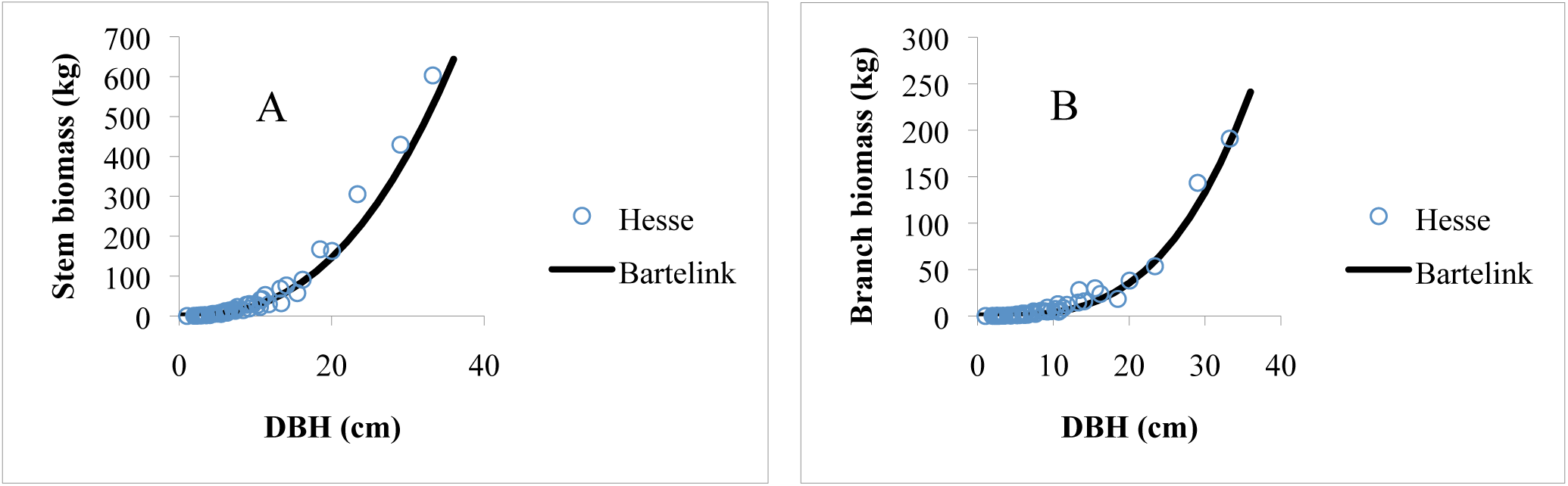
Stem (A) and branch (B) biomass data in relation with tree diameter at breast height (DBH), as observed from the Hesse sample (this study) and from the relation established by Bartelink (1997) which explains 90% of the variation in his sample (Hesse sample here restricted to fit the age range of Bartelink’s study).

The root system contributed less than 20% to tree biomass, this proportion appearing relatively independent of tree age and status, except maybe for the smallest trees of young age classes (Fig. 7). For very young trees, the root/shoot ratio amounted to 0.32 while the mean value for the whole tree sample was 0.23: these values are consistent with those found by other authors in Germany (Bolte & al., 2004) and Central Europe (Konôpka et al., 2010) and with those extracted by Genet & al. (2010) from a literature review. The decrease of the root/shoot ratio observed with increasing age may be the consequence of an ontogenetic drift with plant size and age (Reich, 2002). The fraction of tree biomass included in the root system appeared then relatively independent of the tree status, as already observed by Bolte & al (2004) for beech, unlike what happened for branches.

Coarse roots contributed between 80 and 90% to total root system biomass, the proportion being relatively constant and close to 90% for mature trees (tree age ≥ 60 years). For younger trees, coarse root biomass contribution tended to decrease with decreasing tree size (Fig. 8), together with a higher contribution of small and fine roots, as already found (Le Goff & Ottorini, 2001). The coarse root biomass data obtained in this study for trees of various ages appear consistent with the data obtained by Pellinen (1986) for trees of various dimensions and aged between 100 and 115 years (Fig. 15), but differ from those obtained by Bolte & al. (2004) which appeared well below those of Pellinen. This could be related to the lower root/shoot ratio observed for beech trees in the case of Bolte & al. study – coarse roots biomass representing the major part of the root system biomass – and explained by differences in the environmental conditions of the different sites (Bolte & al., 2004).

**Figure 15.**
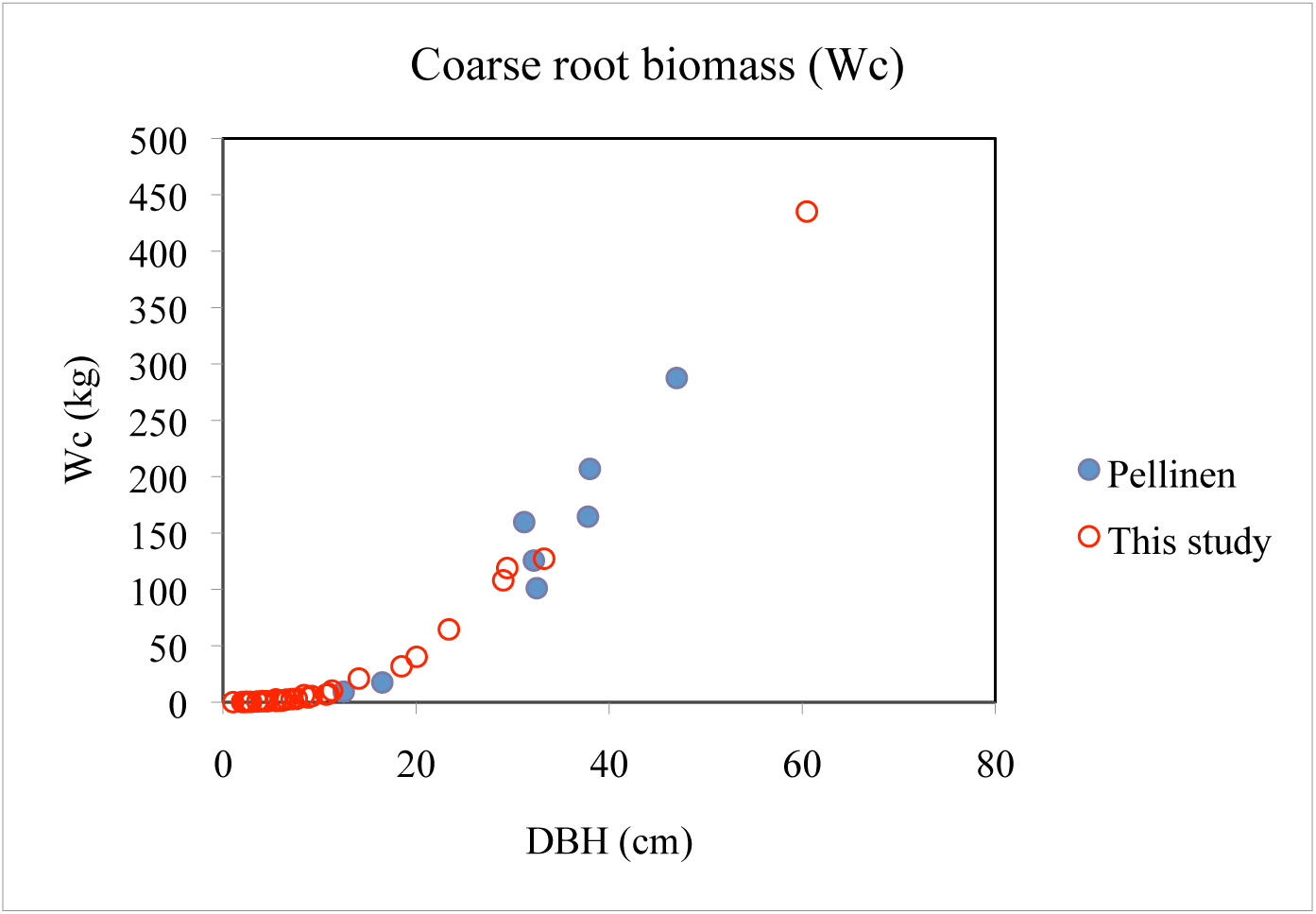
Coarse root biomass data in relation to DBH, for the trees sampled in Hesse forest (this study) compared with those obtained by Pellinen (1986) in Germany.

### 4.4 Biomass increment

#### 4.4.1 Biomass increment models

The sigmoid model described by Eq.11 was successfully fitted for above and belowground components. Moreover, the increment of the different components of the above and belowground biomasses appeared proportional to total aboveground and total belowground biomasses respectively, the increment models differing only by a multiplicative parameter (or allocation coefficient) in each case. Then, 76% of aboveground biomass increment appeared allocated to the stem, compared to 24% to branches, while 95% of belowground biomass increment was allocated to coarse roots compared to 5% to small roots. The biomass increment of aboveground components appeared dependent on forest plot, in relation probably with varying environmental conditions, which was not the case for belowground compartments. However, no effect of varying climatic conditions over sampled years could be detected, in relation maybe with the sampling scheme of the study where forest plots were not sampled every sampled year, which may have led to confused stand and climatic effects.

The biomass increment models fitted (Eq.11 & Eq.13) show that biomass increment depends not only on tree leaf area – as in the basic model used by Hofmeyer & al. (2010) to describe bole volume increment – but also on foliage density and age. Biomass increment increases with tree leaf area, but at a slowing rate as trees are ageing (Fig. 16), while foliar density (*DSF*), which decreases exponentially with age (Eq.15), shows a positive effect on tree biomass increment when it decreases. While the 3-variable model fitted explains more variation than the model with only leaf area as independent variable, there remains a large unexplained variation that could eventually be reduced with a larger sample of trees.

**Figure 16.**
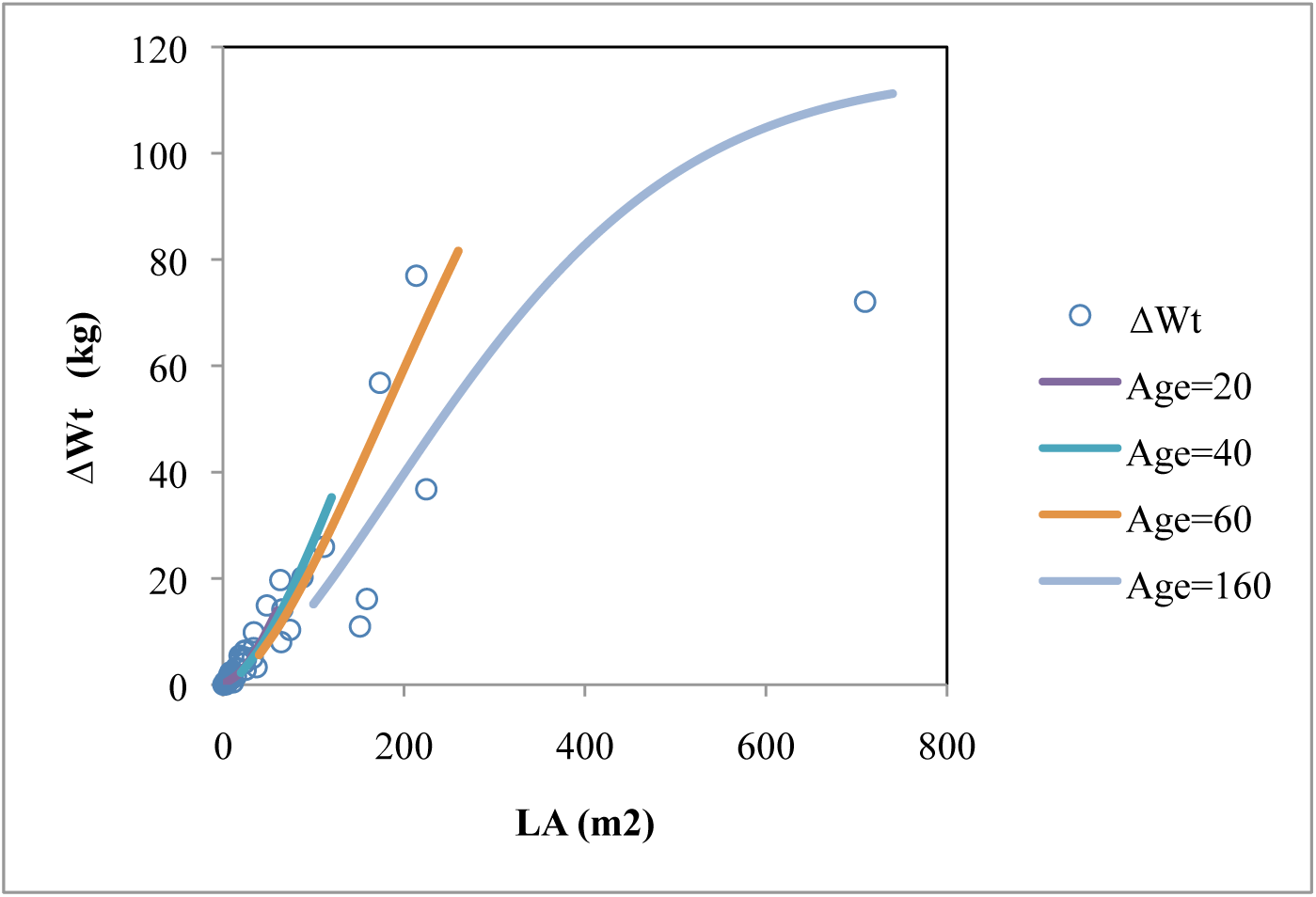
Observed aboveground annual biomass increment data (Δ*Wt*) in relation with tree leaf area (*LA*) and projected values obtained from Eq. 12 for trees of ages covering the sample age range (in Eq. 12, the density of foliage was estimated from the relation established with tree age, that is Eq. 15).

#### 4.4.2 Tree biomass allocation

Relating leaf area and density of foliage to tree age allowed to representing the evolution with age of the allocation of biomass increment to the main tree components (Fig. 11). It appeared that the stem contributed to more than 60% of tree biomass increment – nearly 70% in young ages – whereas branches contributed to a relatively constant fraction close to 20% and roots to less than 20% (only 10% in young ages). These proportions compare relatively well with those obtained on ash (*Fraxinus excelsior* L.) in a study conducted in a nearby region on a sample of trees aged 25 (Le Goff & al. 2004), whereas some variations seemed to occur between ash trees of different competitive status, which was not the case here with beech.

### 4.5 Stem volume increment and growth efficiency of trees

The sigmoid model fitted to biomass increment data was successfully fitted to bole volume increment data (Eq. 16), not surprisingly as stem biomass increment appeared proportional to stem volume increment. Thus, the mean stem wood density^17^ of the beech sample, which appears to be the slope of the linear relation fitted between stem biomass and volume increments, was equal to 0.549 (Fig. 17). This density value is in the range of the observed values for different beech samples in France and other countries in Europe (Nepveu, 1981), and is close to the mean value (0.556) obtained from biomass equations by Genet et al. (2011b).

**Figure 17.**
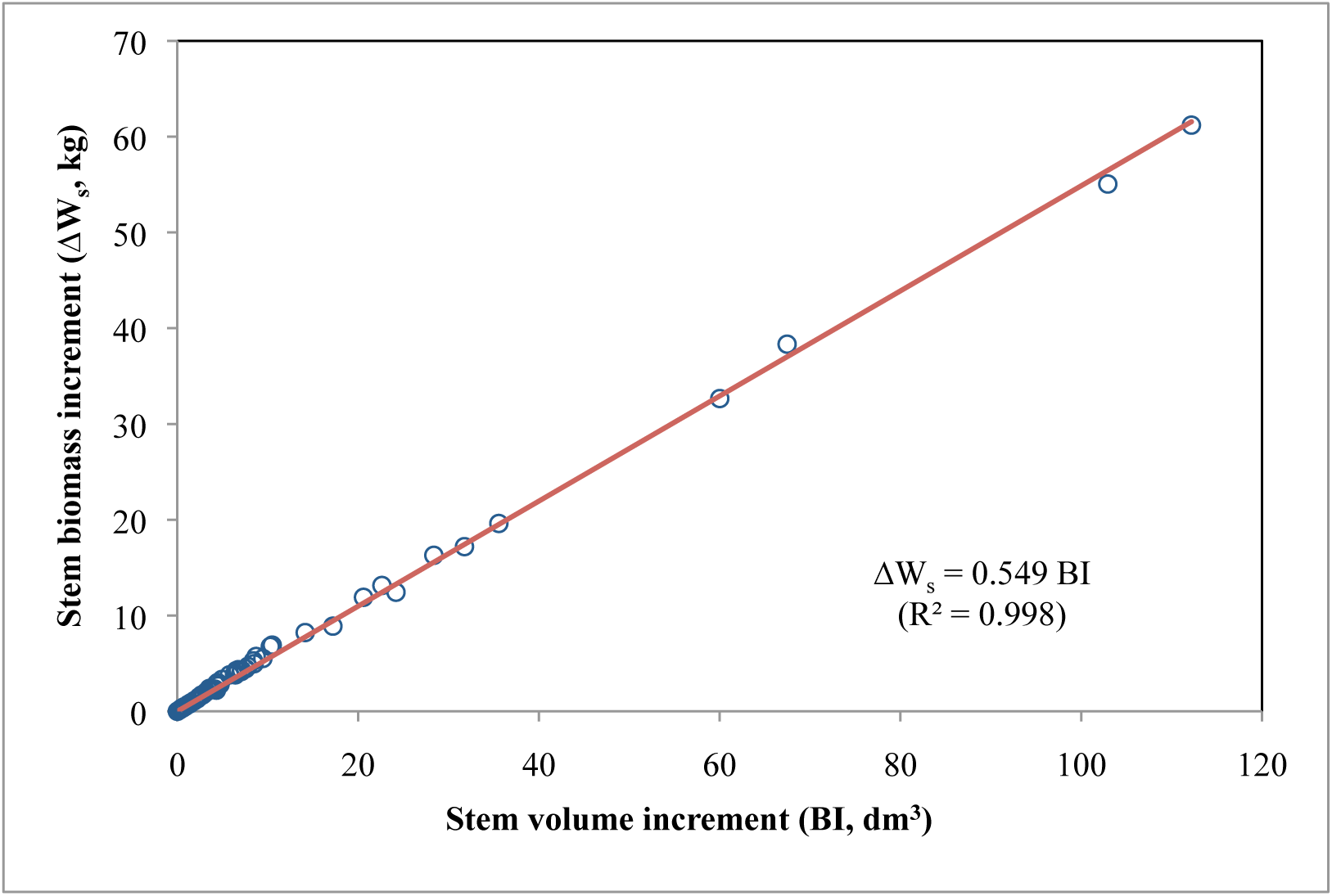
Annual stem biomass increment (Δ*W_s_*) in relation with stem volume increment (*BI*): observed data and linear relation fitted Δ *W_s_* = *ρ_s_ BI*, with *ρ_s_* = 0.549 kg/dm3 (R^2^ = 0.99).

As for biomass increment, a “forest plot” effect was detected for stem volume increment, probably also in relation with environmental conditions (soil and climate).

Stem wood growth efficiency (*GE*), defined here as annual stem volume increment per unit of leaf area, appeared to increase rapidly with age until trees reached the age of 20–30 years (Fig. 13), and then increased more slowly until it began to decrease after the age of about 100 years. This result was obtained after taking account of the variations of tree leaf area (*LA*) and density of foliage (*DSF*) with tree age (Eq. 14 & Eq.15 respectively). The decline of *GE* with age, after culminating at relatively young ages, has been already reported for coniferous species, at tree level, (Kaufmann & Ryan, 1986; Ryan & al., 1997; Day & al, 2001). But other studies failed to reveal such a decrease of *GE* with age when leaf area effects were not taken into account (Seymour & al., 2002; Harper, 2008). The pattern of variation of *GE* with age could be either attributable to variations of productivity per unit of leaf area or to variations of biomass allocation, as trees get older. However, no important variation of biomass allocation with tree age could be observed (Fig. 11), only a small advantage for the stem at an early age. Then, the pattern of variation of *GE* with age could be attributable to an ontogenetic effect (Day & al., 2001, Seymour & al., 2002), the decrease of *GE* at higher ages reflecting probably a less efficient physiological functioning of trees as they get older (Ryan & al., 1997).

As shown by Fig. 18, growth efficiency *GE* depends not only on age, but also on leaf area (*LA*) and density of foliage (*DSF*). Thus, the asymptotic model for *BI* (Eq. 11) predicts also a declining *GE* with increasing tree leaf area (cf Maguire & al, 1998; Seymour & al, 2002), but only above a value of about 300 m^2^ for leaf area, only observed in the oldest trees of the sample. *GE* decreases also, for a given *LA*, with increasing values of the density of foliage, which is the ratio of leaf area on transformed bole area: such a decrease, already observed with ash (Le Goff & al, 1996), a less shade tolerant species than beech, may be related to a less favorable ratio of assimilatory to respiratory processes associated with the increase of foliage area per unit of bole surface area. But, conversely, the decrease of the density of foliage with age, as observed in the sample, contributes to counterbalance the negative effect of increasing age on the growth efficiency of trees. Moreover, there is a relationship between the density of foliage and the crown ratio that shows a minimum for trees moderately crowded (crown ratio of about 0.45): thus, those trees exhibit a better growth efficiency compared to trees less or more crowded, which contrasts with the results obtained with ash (Le Goff & al, 1996).

**Figure 18.**
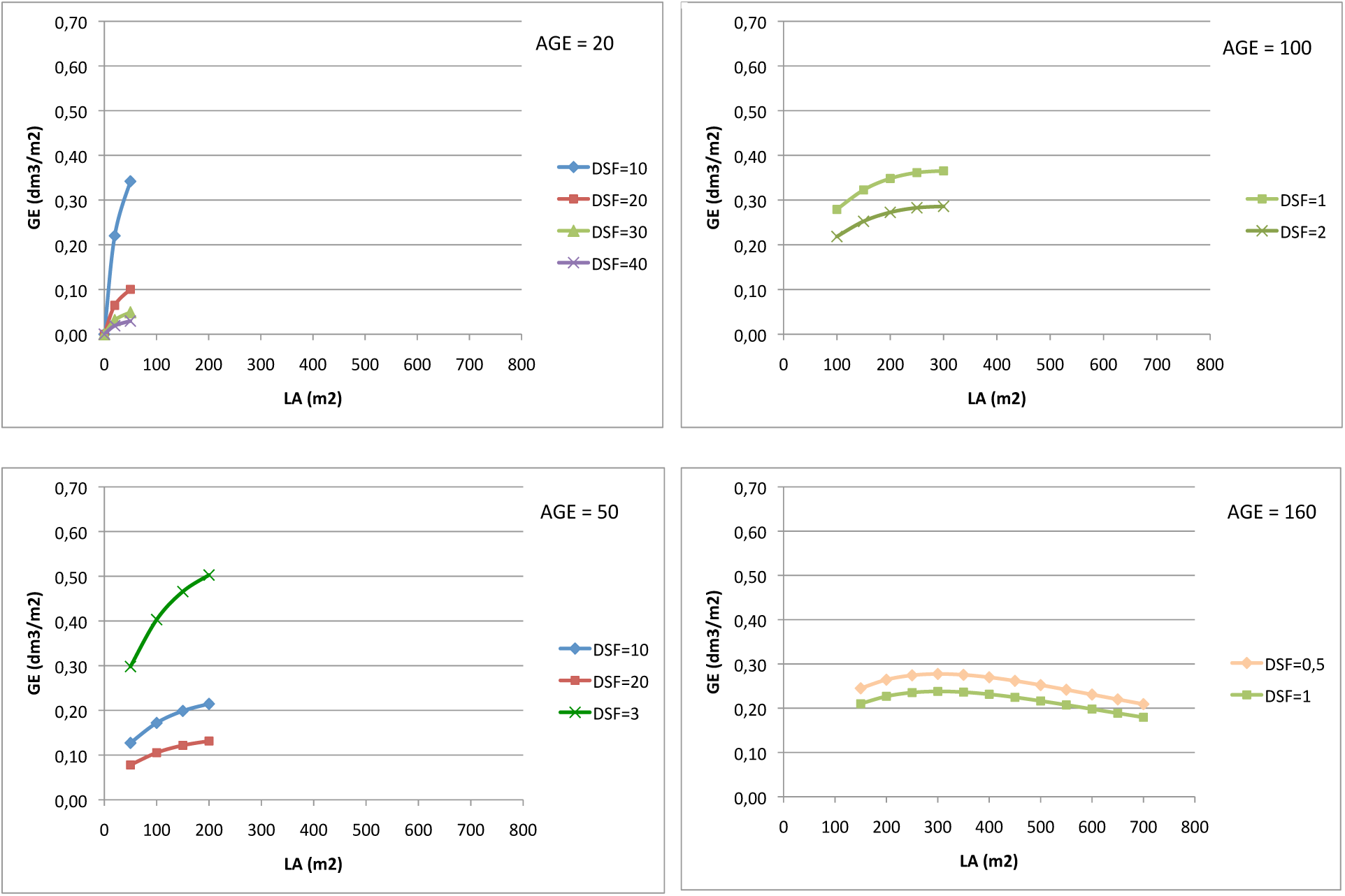
Growth efficiency *(GE*) of beech, in relation with leaf area (*LA*) and density of foliage (*DSF*), for trees of increasing ages (from 20 to 160 years), (the curves represented were restricted to the range of leaf areas and densities of foliage observed according to age in the tree sample)

Then, the most growth efficient beech trees appear to be middle-aged (around 50 years old), dominant with relatively large crowns (leaf area around 200 m^2^) and moderately crowded (crown ratio around 0.45). Such trees exhibit an annual stem volume increment of about 100 dm^3^.

### 4.6 Use of biomass and biomass increment models

The biomass and biomass increment models established for beech in this study allow the estimation of the biomass and carbon stocks and fluxes for the even-aged beech stands of Hesse forest, whatever the age of the stand. Thus, it should help to extend the studies on the ecophysiological functioning of beech stands presently conducted in Hesse forest (Granier et al., 2008) to younger and older stands, and in particular the comparison of the net primary productivity (NPP) of stands estimated from the CO_2_ fluxes with the stand biomass increment (Granier et al., 2000). Moreover, it could help to test the ability of biogeochemical models, like BIOME-BGC, to assess the gross and net primary production of beech stands, as was done by Chiesi & al. (2014) for beech forests in Italy.

The biomass equations established could also be used to analyze the effects of different silvicultural treatments on the biomass and carbon stocks and fluxes of beech stands, using the available stand growth and yield models built in France, that is *Fagacées* (Dhôte & Le Moguedec, 2005) or *SimCAP* (Ottorini & Le Goff, 2006).

The generalized biomass and biomass increment equations established for Hesse forest should however be used with care for beech stands of other regions differing by site conditions, although the models developed for biomass are very similar to the ones developed at a larger scale by Genet et al. (2011). More confident data are still necessary to obtain, in particular for the root biomass compartments, so as to develop more precise biomass and biomass increment equations.

## Acknowledgments

Our thanks go the INRA technicians of the LERFoB research unit at INRA-Nancy, who proceeded to the tree measurements: R. Canta, F. Bordat, G. Maréchal and S. Daviller. Special thanks are addressed to R. Canta who supervised the field and laboratory measurements and developed a specific apparatus to allow the biomass measurements on the large root systems. The support of the ONF section of Sarrebourg (57), greatly appreciated, made possible the felling of the tree samples used in this study. We would like also to thank warmly A. Granier (EEF, INRA-Nancy) for his interest in the biomass studies that we conducted in the Hesse forest over several years, in parallel with the ecophysiological studies that he conducted himself, and for his comments on the first draft of the manuscript.

## Footnotes

1 In this paper, tree diameter is the diameter of the tree at breast height (1.3 m).

2 The definitions of biomass allocation and distribution agree with those used by Reich (2002).

3 Crown base is here defined as the point of the stem where is inserted the first main branch constituting the crown.

4 The crown projection points are the projections of a set of points taken on the periphery of the crown so that its maximum extension in all directions is represented.

5 In this case, only the leaf biomass of trees for year 2001 could not be reconstructed.

6 In this case, 2 or 3 branches were sampled in each of the two crown strata defined, depending on the tree sample.

7 For the samples with an irregular shape, a correction was applied to the calculated sample section area by comparing it to the area of the same section scanned.

8 In this paper, as recommended by Sileshi (2014), it will not be referred to “allometric models” to design the biomass equations used, as they don’t follow the typical power-law function.

9 The relative crown length (*RCL*) of a tree was defined as the ratio of crown length (distance from crown base to stem apex) to total tree height (*Ht*).

10 Growth efficiency, defined as stem volume increment per unit of leaf area, presents a peaking pattern when plotted against tree leaf area (Hofmeyer & al, 2010; Seymour and Kenefic, 2002).

11 Small and fine roots data of tree # 35 were dropped from the belowground data file, as they appeared abnormally low.

12 In fact, the measured root biomasses of tree # 35 were more probably underestimated due to the difficulty to estimate the biomass of the missing parts of the root system after soil extraction.

13 The root-shoot ratio of the sampled trees was calculated as the ratio of total belowground biomass on total aboveground biomass of trees (Reich, 2002).

14 Foliar density is here defined as the ratio *LA/BS*^1.5^ where *LA* is tree leaf area and *BS* is stem surface area (a similar ratio where leaf biomass replaced leaf area was used in a preceding study: Le Goff & Ottorini, 1996).

15 *DSF* values obtained from Eq. 14 were multiplied by the correction factor 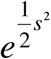

16 Fine roots biomass increment was not evaluated in this study, as the main part of it is due to fine root turnover whose quantification would have necessitated specific studies (Le Goff & Ottorini, 2001).

17 Wood density, or wood specific gravity (*ρ_s_*), is here defined as the ratio of dry weight on green volume.

## References

Bartelink H. H. (1997) Allometric relationships for biomass and leaf area of beech (*Fagus sylvatica* L). Ann. Sci. For. 54: 39–50.

Bolker B. M. (2008) Ecological Models and Data in R. Princeton University Press, 41 William Street, Princeton, New Jersey 08540, USA.

Bolte A., Rahmann T., Kuhr M., Pogoda P., Murach D., Gadow K. v. (2004) Relationships between tree dimension and coarse root biomass in mixed stands of European beech (Fagus sylvatica L.) and Norway spruce (Picea abies [L.] Karst.). Plant and Soil, 264: 1–11.

Bond-Lamberty B., Wang C., Gower S.T. (2002) Aboveground and belowground biomass and sapwood area allometric equations for six boreal tree species of northern Manitoba. Can. J. For. Res. 32: 1441–1450.

Bouriaud O., Breda N., Le Moguedec G., Nepveu G. (2004) Modelling variability of wood density in beech as affected by ring age, radial growth and climate. Trees, Structure and Function, vol. 18 (3): 264–276.

Breda N. (1999) L’indice foliaire des couverts forestiers: mesure, variability et role fonctionnel. Rev. For. Fr. 51(2): 135–150.

Breda N., Granier A. (1996) Intra- and inter-annual variations of transpiration, leaf area index and radial growth of a sessile oak stand (*Quercus petraea*). Ann. Sc. For., vol. 53 : 521–536.

Cannell M.G.R (1989) Physiological basis of wood production: a review. Scand. J. For. Res., 4: 459–490.

Chiesi M., Maselli F., Chirici G., Corona P., Lombardi F., Tognetti R., Marchetti M. (2014) Assessing most relevant factors to simulate current annual increments of beech forests in Italy. iForest (2014) 7: 115–122.

DeRose R. J., Seymour R.S. (2009) The effect of site quality on growth efficiency of upper crown class Picea rubens and Abies balsamea in Maine, USA. Can J For Res 39: 777–784.

Dhôte J.-F., Le Moguedec G. (2005) Présentation du modèle Fagacées. Nancy : LERFoB, UMR 1092 INRA-ENGREF (Document interne).

Flewelling J.W., Pienaar L.V. (1981) Multiplicative regression with lognormal errors. For. Sci. 27 : 281–289.

Genet A., Wernsdörfer H., Jonard M., Pretzsch H., Rauch M., Ponette Q., Nys C., Legout A., Ranger J., Vallet P., Saint-André L. (2011a) Ontogeny partly explains the apparent heterogeneity of published biomass equations for *Fagus sylvatica* in central Europe. For. Ecol. Manage. 261:1188–1202.

Genet A., Wernsdörfer H., Mothe F., Bock J., Ponette Q., Jonard M., Nys C., Legout A., Ranger J., Vallet P., Saint-André L. (2011b) Des modèles robustes et génériques de biomasse. Exemple du Hêtre. Rev For Fr 63:179–190.

Genet H., Bréda N., Dufrêne E. (2010) Age-related variation in carbon allocation at tree and stand scales in beech (*Fagus sylvatica* L.) and sessile oak (*Quercus petraea* (Matt.) Liebl.) using a chronosequence approach. Tree Physiology 30, 177–192.

Granier A., Ceschia E., Damesin C., Dufrêne E., Epron, D., Gross P., Lebaube S., Le Dantec V., Le Goff N., Lemoine D., Lucot E., Ottorini J.-M., Pontailler J.Y., Saugier B. (2000) The carbon balance of a young beech forest. Funct. Ecol. 14: 312–325.

Granier A., Brédat N., Longdoz B., Gross P., Ngao J. (2008) Ten years of fluxes and stand growth in a young beech forest at Hesse. North-eastern France. Ann. For. Sci. 64: 704p1–704p13.

Harper G. (2008) Quantifying branch, crown and bole development in *Populus tremuloïdes* Michx. from north-eastern British Columbia. For. Ecol. Manage. 255: 2286–2296.

Hofmeyer P. V., Seymour R. S., Kenefic L. S. (2010) Production ecology of *Thuya occidentalis*. Can J For Res 40: 1155–1164.

Kaufmann M. R., Ryan M. G. (1986) Physiographic, stand, and environmental effects on individual tree growth and growth efficiency in subalpine forests. Tree Physiology 2, 47–59.

Konôpka B., Pajtik J., Moravcik M., Lukac M. (2010) Biomass partitioning and growth efficiency in four naturally regenerated forest tree species. Basic and Applied Ecology 11: 234–243.

Lebaube S., Le Goff N., Ottorini J.-M., Granier A. (2000) Carbon balance and tree growth in a *Fagus sylvatica* stand. Ann For Sci 57 : 49–61.

Le Dantec V., Dufrene E., Saugier B. (2000) Interannual and spatial variation in maximum leaf area index of temperate deciduous stands. For. Ecol. Manage. 134:71–81.

Le Goff N., Ottorini J.-M. (1996) Leaf development and stem growth of ash (Fraxinus excelsior L.) as affected by tree competitive status. Journal of Applied Ecology 33: 793–802.

Le Goff N., Ottorini J.-M. (1998) Biomasses aériennes et racinaires et accroissements annuels en biomasse dans le dispositive écophysiologique de la forêt de Hesse. Rapport scientifique annuel, Contrat ONF-INRA “Croissance du Hêtre sur le Plateau lorrain”, 29pp.

Le Goff N., Ottorini J.-M. (1999) Effets des éclaircies sur la croissance du hêtre. Interaction avec les facteurs climatiques. R.F.F., LI-2, 355–364.

Le Goff N., Ottorini J.-M. (2001) Root biomass and biomass increment in a beech (Fagus sylvatica L.) stand in North-East France. Ann For Sci 58: 1–13.

Le Goff N., Granier A., Ottorini J.-M., Peiffer M. (2004) Biomass increment and carbon balance of ash (*Fraxinus excelsior*) trees in an experimental stand in northeastern France. Ann. For. Sci. 61 : 1–12.

Maguire D. A., Brissette J. C., Gu L. (1998) Crown structure and growth efficiency of red spruce in even-aged, mixed-species stands in Maine. Can. J. For. Res. 28: 1233–1240.

McElligott K. M., Bragg D. C. (2013) Deriving biomass models for Small-diameter Loblolly Pine on the Crossett Experimental Forest. Journal of the Arkensas Academy of Science, Vol. 67 : 94–101.

Nepveu G. (1981) Propriétés du bois de Hêtre. *In* : Le Hêtre, Monographie INRA, Paris, 1981, pp 377–387.

Oliver C.D., Larson B.C. (1996) Forest stand dynamics. John Wiley and Sons, Inc., New York, USA.

Ottorini J.-M., Le Goff N. (1999) Aspects quantitatifs et qualitatifs de la biomasse. Rapport scientifique final (3ième année), Convention de recherche ONF-INRA “Etude de la croissance du hêtre sur le Plateau lorrain”, Juillet 1999, 18 p.

Ottorini, J.-M., Le Goff, N. (2006). *SimCAP*, Simulation et intégration des connaissances : données expérimentales et simulées de la croissance du Frêne et du Hêtre. Conseil Scientifique LERFoB 2006, 14 mars 2006, ENGREF, Nancy (document “PowerPoint”), 21 pp.

Parresol, B. R. (2001). Additivity of nonlinear biomass equations. Can. J. For. Res. 31: 865–878.

Pauwels, D. (2001). Le Vertex, une nouvelle génèration de dendromètres multi-usages. Note technique forestière de Gembloux n°1, mai 2001. Faculté universitaire des sciences agronomiques, Gembloux, Belgique.

Pellinen P. (1986) Biomasseuntersuchungen im Kalkbuchenwald, Dissertation Universität Göttingen, Germany, 134 p.

Pineiro, G., Perelman S., Guerschman J. P., Paruelo J. M. (2008) How to evaluate models : Observed vs. Predicted or predicted vs. Observed ?. Ecological Modelling, 216, 316–322.

R Development Core Team (2009) R: A Language and Environment for Statistical Computing. R Foundation for Statistical Computing, Vienna, Austria. ISBN 3900051-070

Reich P. B. (2002) Root-shoot relations: optimality in acclimation and adaptation or the “Emperor’s New Clothes”? In: Plant Roots, The Hidden Half, 3rd Ed., Marcel Dekker, Inc., New York.

Repola J. (2008) Biomass equations for Birch in Finland. Silva Fennica 43 : 605–623.

Repola J. (2009) Biomass equations for Scots pine and Norway spruce in Finland. Silva Fennica 43 : 625–647.

Ritz C., Streibig J.C. (2008) Nonlinear Regression with R. Springer Science+Business Media, LLC, 233 Spring Street, New York, NY 10013, USA.

Ryan M.G., Binkley D., Fownes J.H. (1997) Age-related decline in forest productivity: Patterns and processes. Adv. Ecol. Res. 27: 213–256.

Seymour R.S., Kenefic L.S. (2002) Influence of age on growth efficiency of *Tsuga canadensis* and *Picea rubens* trees in mixed-species, multiaged northern conifer stands. Can J For Res 32: 2032–2042.

Shaiek O., Loustau D., Trichet P., Meredieu C., Bachtobji B., Garchi S., EL Aouni M.H. (2011) Generalized biomass equations for the main aboveground biomass components of maritime pine across contrasting environments. Ann. For. Sci. 68:443–452.

Sileshi G. W. (2014) A critical review of forest biomass estimation models, common mistakes and corrective measures. For. Ecol. Manage. 329: 237–254.

Velleman P.F. (2011) Data Desk 6.3, Data Description Inc., P.O. Box 4555, Ithaca, NY 14852, USA.

Zeng W.S., Zhang H.R., Tang S.Z. (2011) Using the dummy variable model approach to construct compatible single-tree biomass equations at different scales — a case study for Masson pine (*Pinus massoniana*) in southern China. Can J For Res 41: 1547–1554.

Zheng C., Mason E.G., Jia L., Wei S., Sun C., Duan J. (2015) A single-tree additive biomass model of *Quercus variabilis* Blume forests in North China. Trees 29: 705–716.

